# The copy-number and varied strengths of MELT motifs in Spc105 balance the strength and responsiveness the Spindle Assembly Checkpoint

**DOI:** 10.1101/2020.01.07.897876

**Authors:** Babhrubahan Roy, Simon JY Han, Adrienne N. Fontan, Ajit P. Joglekar

## Abstract

The Spindle Assembly Checkpoint (SAC) maintains genome stability while enabling timely anaphase onset. To maintain genome stability, the SAC must be strong so that it delays cell division even if one chromosome is unattached, but for timely anaphase onset, it must be responsive to silencing mechanisms. How it meets these potentially antagonistic requirements is unclear. Here we show that the balance between SAC strength and responsiveness is determined by the number of ‘MELT’ motifs in the kinetochore protein Spc105/KNL1 and their Bub3-Bub1 binding affinities. Spc105/KNL1 must contain many strong MELT motifs to prevent chromosome missegregation, but not too many, because this delays SAC silencing and anaphase onset. We demonstrate this by constructing a Spc105 variant that trades SAC responsiveness for significantly improved chromosome segregation accuracy. We propose that the necessity of balancing SAC strength with responsiveness drives the evolutionary trend of MELT motif number amplification and degeneration of their functionally optimal amino acid sequence.

## Introduction

To achieve accurate chromosome segregation, the dividing cell executes three processes prior to anaphase onset: (1) assembly of a bipolar spindle, (2) capture of unattached chromosomes by the spindle, and (3) bipolar attachment of each chromosome to microtubules emanating from opposite spindle poles. This requires some minimum amount of time for proper execution. Moreover, the time required for chromosome capture and biorientation is not completely predictable; stochastic events can introduce significant delays in their completion. To provide a minimum amount of time to the cell for spindle formation, and to prolong cell division as necessary to ensure chromosome attachment to the spindle, the eukaryotic cell uses a signaling mechanism known as the ‘Spindle Assembly Checkpoint’ (SAC). The SAC is activated by a signaling cascade that operates from unattached kinetochores, protein structures that mediate chromosome attachment to the spindle. It produces an anaphase-inhibitory signal that delays cell division (Musacchio, 2015). To ensure both accurate chromosome segregation and timely anaphase onset, the dividing cell must ensure that SAC signaling is strong, but also responsive to silencing mechanisms. Whether such balanced SAC operation is important, and if so, how it is achieved remains poorly understood.

A key determinant of the strength of the SAC is the conserved kinetochore protein Spc105, which provides the physical scaffold for SAC signaling. Spc105 and its homologs possess a large, unstructured phosphodomain containing short sequence repeats commonly referred to as ‘MELT’ motifs because of their consensus sequence in yeast and humans. Phosphorylation of the MELT motif by the conserved Mps1 kinase turns on the SAC signaling cascade and results in the generation of the anaphase-inhibitory signal (London et al., 2012; Meadows et al., 2011a; Vleugel et al., 2013) (Fig. 1A). Even just one MELT motif can delay anaphase when phosphorylated in yeast and human cells (Aravamudhan et al., 2016; Chen et al., 2019; Krenn et al., 2014). Therefore, the number of MELT motifs per Spc105 molecule is a crucial determinant of the strength of SAC signaling, and many MELT motifs per Spc105 are expected to endow it, and by extension the kinetochore, with a larger signaling capacity. Indeed, Spc105 and its homologs typically contain many MELT motifs, e.g. six in budding yeast and nineteen in humans (Tromer et al., 2015), and the large MELT motif number is thought to be essential for implementing a strong SAC (Chen et al., 2019; Krenn et al., 2014; Vleugel et al., 2015; Vleugel et al., 2013; Zhang et al., 2014). However, when budding yeast and human cells are treated with high doses of the microtubule poison nocodazole, only a small fraction of MELT motifs, ∼ 20 and ∼ 35% respectively, engage in SAC signaling, and even fewer MELT motifs are sufficient for arresting cell division (Aravamudhan et al., 2016; Vleugel et al., 2015; Vleugel et al., 2013; Zhang et al., 2014). These observations bring into question the simple model that the large number of MELT motifs achieve a correspondingly large amplification of the signaling capacity of Spc105.

**Figure 1.**
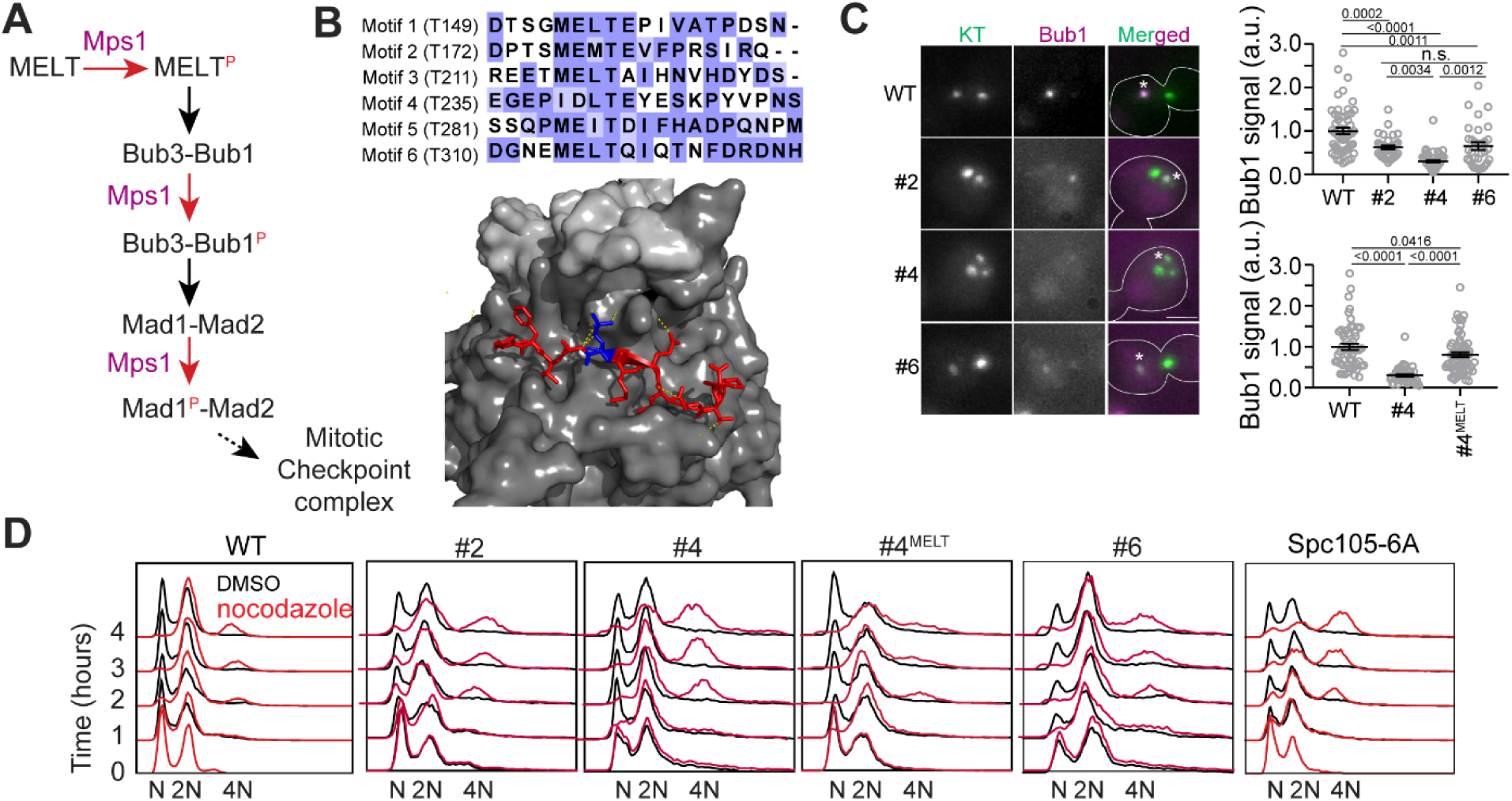
Only one MELT motif per Spc105 is sufficient for SAC signaling in nocodazole-treated cells. **A** A simplified schematic of the signaling cascade of the SAC in budding yeast (Mad3/BubR1, Cdc20, etc. are not shown). **B** Top: amino acid sequence alignment for the six MELT motifs in the *Saccharomyces cerevisiae* Spc105. Bottom: Crystal structure of the MELpT-Bub3-Bub1 complex (adapted from PDB 4BL0 and 2I3S). Note that the phosphothreonine (shown in blue, inter-chain hydrogen bonds highlighted by dashed lines) buried in a grove on Bub3 surface. **C** Micrographs display representative images of nocodazole-treated yeast cells with fluorescently labeled kinetochores and Bub1-mCherry. Asterisks mark Bub1-mCherry colocalized with the unattached kinetochore clusters. Bar∼3.2µm. Scatterplots show the quantification of Bub1-mCherry fluorescence in wild-type (WT) and mutant strains (labeled by the number designation of the active MELT motif). Fluorescence values have been normalized to the average Bub1-mCherry fluorescence in wild-type cells. (mean+s.e.m. Top: n=55, 33, 51 and 32 for WT, #2, #4 and #6 respectively, accumulated from two technical replicates. Bottom: n=55, 92 and 68 for WT, #2 and #6 respectively, pooled from two technical replicates. n.s. Not significant. **D** Quantification of DNA content in cells treated with nocodazole using flow cytometry (one of two biological replicates displayed).

The amino acid sequence of a MELT motif directly determines its SAC signaling activity, because it directly binds to the Bub3-Bub1 complex (Fig. 1B) (Primorac et al., 2013). However, bioinformatics analyses show that in many eukaryotes, the MELT motif number as well as sequence is evolving rapidly due to: (a) duplication and loss of existing MELT motifs, and (b) sequence divergence of a subset of the MELT motifs from the consensus amino acid sequence (Tromer et al., 2015). In human and yeast cells, sequence divergence has created many MELT motifs with a weak ability to bind Bub3-Bub1, and hence a significantly lower signaling activity (Primorac et al., 2013; Vleugel et al., 2015; Vleugel et al., 2013). Why Spc105 contains a large number of MELT motifs, but then uses only a small fraction under certain experimental conditions, and why the sequence of many MELT motifs has diverged from the functionally optimal sequence remains unknown. Here we show that the number of MELT motifs in Spc105 and their sequences strongly influence the strength of SAC signaling and its responsiveness to silencing mechanisms.

## Results

### In nocodazole-treated budding yeast cells, Spc105 with just one MELT motif is sufficient to effect a prolonged mitotic arrest

Three of the six motifs in Spc105 possess the consensus amino acid sequence (‘MELT’), and are likely to display higher Bub3-Bub1 binding affinity than the remaining three motifs possessing divergent amino acid sequences (London et al., 2012). We wanted to verify this expectation and also determine the minimum number of MELT motifs per Spc105 required to maintain a prolonged mitotic block.

We created three different yeast strains each expressing a Spc105 mutant wherein either motif #2, #4, or #6 is the only phosphorylatable, active motif (numbering follows Fig. 1B). To study SAC signaling in these strains, we destroyed the mitotic spindle using the microtubule poison nocodazole to activate the SAC. To measure the signaling activity of each motif, we quantified the recruitment of Bub1-mCherry by unattached kinetochore clusters in nocodazole-treated cells (Fig. 1C, Methods). To assess the SAC strength in these strains, we quantified the ploidy of cell populations over four hours of nocodazole treatment (Fig. 1D). Motif #2 and #6 supported robust SAC signaling as judged by the ability of the mutant cells to maintain 2N ploidy over the 4-hour long duration of the nocodazole-induced mitotic arrest despite recruiting ∼50% less Bub1 compared to wild-type Spc105 (the leftmost graph in Fig. 1D, the small fraction of 4N ploidy cells emerging after > 2 hours of nocodazole treatment represents cells that slipped out of the SAC arrest). Motif #4 recruited significantly less Bub1-mCherry than the other two, which is consistent with its weaker affinity for binding recombinant Bub3-Bub1 complex *in vitro* (Overlack et al., 2015). The SAC was also weaker in this mutant as evidenced by the emergence of a considerable fraction of tetraploid cells starting at 2 hours post treatment. When the divergent amino acid sequence of this motif (‘IDLT’) was replaced with the consensus sequence (‘MELT’), this improved both Bub1 recruitment and SAC strength. Thus, the amino acid sequence of each MELT motif determines is ability to recruit Bub3-Bub1 and its contribution to SAC signaling in unattached kinetochores (Fig. 1C-D). Furthermore, just one strong MELT motif per Spc105 is sufficient to induce a prolonged mitotic arrest upon spindle depolymerization by nocodazole.

### Multiple MELT motifs per Spc105 are essential when chromosome attachment to the spindle and biorientation is challenged

The use of nocodazole for quantifying SAC strength has a significant drawback: it only reveals SAC strength in the presence of a large number of unattached kinetochores. Furthermore, the continued presence of nocodazole in growth media prevents spindle assembly and keeps most, if not all, kinetochores unattached. Consequently, even if the signaling capacity of individual kinetochores is impaired, the cumulative output of many signaling kinetochores may still impose a prolonged mitotic arrest under steady state signaling condition (in regard to the number of unattached kinetochores). However, this situation does not occur during the course of a normal cell division. As chromosomes attach to the newly formed spindle, the number of unattached kinetochores decreases rapidly until only a few or just one kinetochore remains unattached. In this dynamic situation, the SAC must reliably delay anaphase irrespective of the number of unattached kinetochores. Therefore, we explored the effect of individual MELT motifs on SAC performance in cells containing small numbers of unattached kinetochores.

We first established conditions to create small numbers of unattached kinetochores using low doses of the microtubule-destabilizing drug benomyl. Unlike nocodazole treatment, which destroys the spindle, low dose of benomyl increases microtubule dynamicity and therefore challenges spindle formation and chromosome biorientation. Wild-type cells still grow under this condition, but any genetic defects in SAC signaling or chromosome biorientation cause poor growth and can result in inviability. We hypothesized that cells grown in the presence of benomyl will more frequently contain small numbers of unattached kinetochore, and therefore would require a strong SAC for proliferation. To test this, we followed chromosome biorientation by marking the centromere of chromosome IV with a TetO array (Fig. 2A left). In normal media, yeast cells released from a G1 arrest bioriented most of their chromosomes in ∼ 45 min and completed anaphase by 75 minutes (biorientation). However, in media containing low doses of benomyl both events were significantly delayed. In most cells, chromosome IV had not achieved biorientation even after 105 minutes after release from a G1-arrest (the duration of the experiment was limited to 105 minutes, because cells not affected by the treatment would enter the next cell cycle after this point complicating analysis). In a small fraction of cells, chromosome IV was displaced from the spindle and was likely unattached. To confirm this, we imaged yeast cells expressing Spc105^222::GFP^ (GFP inserted after the 222^nd^ residue in Spc105) and Bub1-mCherry to assess the attachment state of all the chromosomes under the same conditions. A large fraction of cells contained a significantly smaller number of unattached kinetochores indicated by Bub1-mCherry recruitment, compared to nocodazole-treated cells (5 +/-4 unattached chromosomes per cell in contrast to ∼ 9 unattached chromosomes in nocodazole-treated cells (Aravamudhan et al., 2016), Fig. 2A right). Unattached kinetochores persisted 105 minutes after the introduction of benomyl (Fig. 2A right).

**Figure 2.**
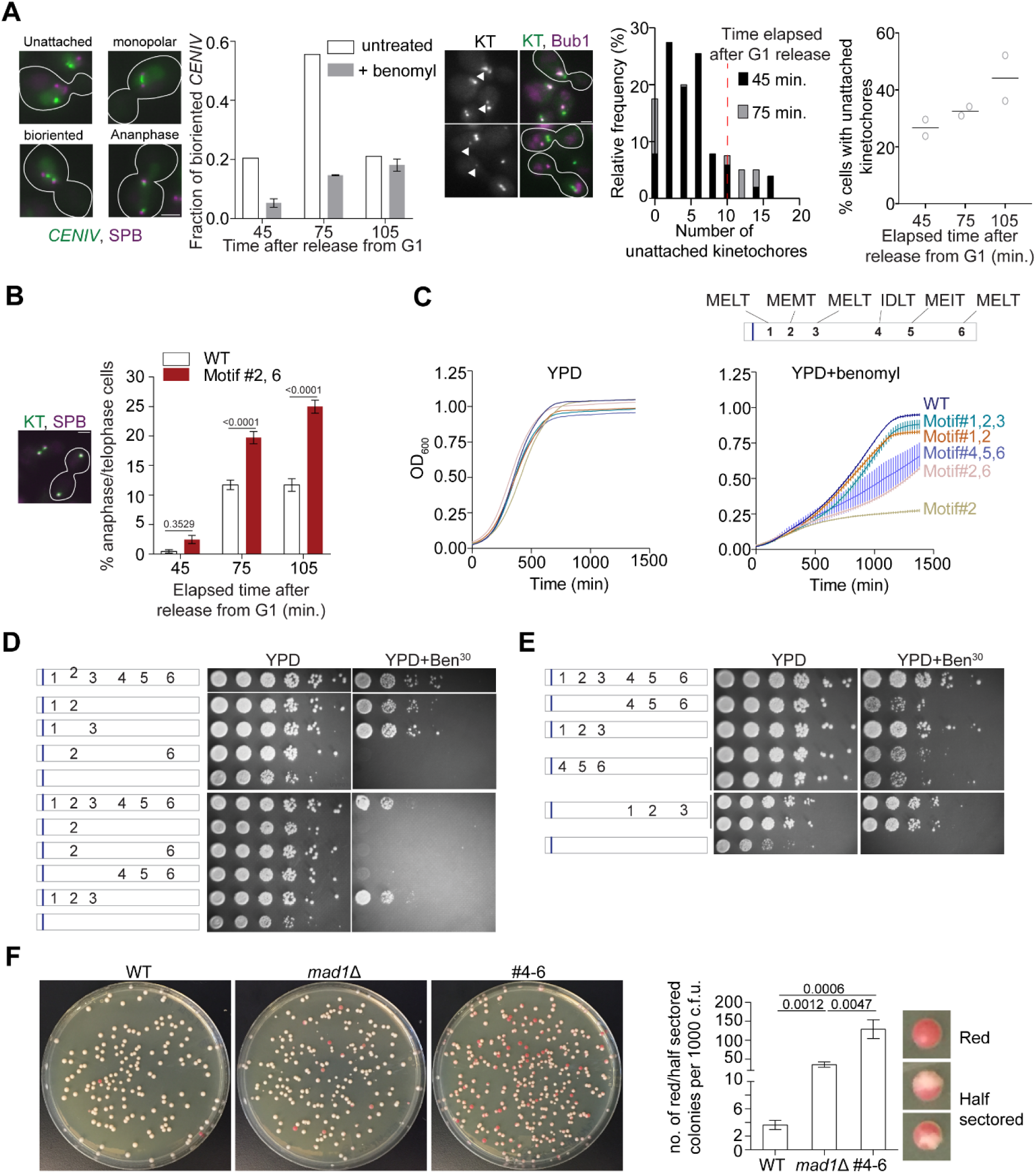
More than one high affinity MELT motifs are necessary to minimize chromosome loss. **A** Effects of benomyl on chromosome biorientation. Left: Kinetics of chromosome IV biorientation visualized using a centromere-proximal TetO array. Micrographs depict representative images of cells showing fluorescently labeled TetO array and Spindle Pole Bodies (SPB). Scale bar∼3.2µm. Second from the left: Quantification of the fraction of cells with bioriented chromosome IV. Note that the reduced fraction of untreated cells with bioriented chromosomes 105 minutes after release from G1 is because most cells complete anaphase by this time. (n = 259, 252 and 398 at 45, 75 and 105 min respectively in normal media, one repeat. n= 855, 816 and 626 respectively at 45, 75 and 105 min in benomyl-containing media from three technical repeats). Middle: Representative micrographs of cells grown in benomyl media containing unattached kinetochores recruiting Bub1. Arrow heads indicate the unattached kinetochores with Bub1 localizations. Scale bar ∼3.2µm. Right: Quantification of the number and frequency distribution of unattached kinetochores in yeast cells growing in media containing benomyl. The unattached kinetochore number was estimated by comparing the Spc105-GFP fluorescence from the unattached kinetochore cluster (marked by Bub1-mCherry recruitment) with the total fluorescence from the kinetochores along the spindle axis. (n= 686, 501 and 543 at 45, 75 and 105 min respectively, pooled from two technical repeats). Red dashed line in relative frequency plot indicates the average number of unattached kinetochores observed after 90min nocodazole treatment of wild type yeast cell (Aravamudhan et al., 2016). **B** Left: Micrograph displays representative cells in metaphase and anaphase in benomyl-containing media. Scale bar ∼3.2µm. Right: Quantification of the number of anaphase cells observed after the indicated time following release from a G1 arrest for the indicated strains. (mean+s.e.m, n=1029, 889 and 992 at 45, 75 and 105 min respectively for WT cells, pooled from three experimental repeats. n=890, 864 and 812 at 45, 75 and 105min time points respectively for #2,6, pooled from three technical replicates). **C** Top: Schematic of the Spc105 phosphodomain with the amino acid sequence indicated at the top. Bottom: Quantification of the evolution of cell density of the indicated strains in rich media (left) and media-containing benomyl (right). **D-E** Assessment of the sensitivity of yeast strains to benomyl using the spotting assay. Schematics on the left show the active motif number and its position. The blue bar represents the ‘basic patch’ in Spc105, which promotes PP1 recruitment. The photographs on the right show the results of spotting a serial dilution of yeast cells on rich media (YPD) and media containing benomyl (20 or 30 μg/ml). **F** Assessment of chromosome loss by colony sectoring assay. Left: Images of WT, *mad1*Δ and #4-6 colonies grown in YPD plates. Right: Bar graph shows the rate of loss of *SUP11* containing chromosome fragment measured as the number of red/half red colonies (example shown on the right) for every 1000 colonies plated. N= 5081, 4406 and 4965 for WT, *mad1*Δ and #4-6 respectively, pooled from at least 6 technical repeats.

We next examined the effects of benomyl on a strain expressing a mutant Spc105 with only two phosphorylatable motifs: #2 and #6, to test whether the SAC might be weaker under this condition. Each of the two MELT motifs by itself supports robust SAC signaling in nocodazole-treated cells (Fig. 1D). 75 minutes after release from the G1 arrest into media containing benomyl more mutant cells were in anaphase compared to wild-type cells, as evidenced by elongated spindles and sister kinetochores segregated in the mother and bud (Fig. 2B). It should be noted that most cells are expected to contain unattached kinetochores at this time point. Therefore, these observations together imply that many cells expressing Spc105 with motif #2 and #6 are unable prevent anaphase onset even though they contain small numbers of unattached kinetochores. Accordingly, many cells expressing the mutant Spc105 displayed elongated spindles with lagging/unaligned kinetochores (Fig. S1A).

If cells expressing a Spc105 mutant mis-segregate their chromosomes more frequently than wild-type cells as a result of weak SAC signaling, the population should display more frequent cell death, and hence reduced growth rates. To test this, we quantified the growth of yeast strains expressing Spc105 mutants carrying different combinations of MELT motifs on normal media and media containing benomyl by quantifying absorbance at 600 nm wavelength (OD_600_, Fig. 2C). In normal media, the mutant strains showed relatively minor differences in their growth rate and the peak cell density (due to nutrient depletion) compared to the wild-type strain (Fig. 2C, top left). However, stark differences in growth rate and maximal cell density were apparent in media containing benomyl (Fig. 2C, top right; for reference a schematic of the MELT motif position and sequences is also included at the bottom of Fig. 2C and in Fig. S1B). Cells expressing Spc105 mutants with either motif #2 alone or motif #2 and #6 together grew at slower rates and reached significantly lower cell densities than wild-type cells. Spc105 mutants containing either the first two or three MELT motifs (two of which possess the consensus amino acid sequence, and hence are expected to be strong), supported growth in benomyl-containing media. Interestingly, the Spc105 mutant with only the last three MELT motifs active grew more slowly and reached lower maximal cell density. Both results are indicative of a weaker SAC. They suggest that in cells containing a small number of unattached kinetochores at least two, strong MELT motifs are necessary for cell survival; the presence of all six MELT motifs in Spc105 ensures maximal population growth. We confirmed these quantitative findings by spotting serial dilutions of yeast cultures on benomyl media (Fig. 2D).

### MELT motif position, their role in Sgo1 recruitment, or pleiotropic effects of benomyl do not explain the correlations between MELT motif mutations and benomyl sensitivity

One possible explanation for the observed variations in benomyl sensitivity of the mutant strains is that the MELT motifs located distally in the Spc105 phosphodomain are situated further away from Mps1 bound to the CH-domains of the Ndc80 complex, and they are therefore less efficiently phosphorylated when compared to the MELT motifs located anteriorly in the phosphodomain (Aravamudhan et al., 2015; Hiruma et al., 2015; Ji et al., 2015; Kemmler et al., 2009). However, even when we swapped the positions of the first three and last three MELT motifs, this did not change the benomyl sensitivities of the two strains (Fig 2E). The lack of correlation persisted even in Spc105 mutants containing only one phosphorylatable MELT motif (see Fig. S1C-D). Thus, the amino acid sequence, and not the position of MELT motifs in the phosphodomain, is the sole determinant of their SAC signaling activity.

In these assays, the differential recruitment of the centromeric protein Sgo1, a factor that is crucial for chromosome biorientation and error correction, can also result in differential benomyl sensitivities (Kawashima et al., 2010a; Kawashima et al., 2010b). Because Bub1 kinase activity promotes the centromeric recruitment of Sgo1, reduced Bub1 recruitment by the Spc105 mutants will reduce centromeric Sgo1 and thus promote chromosome missegregation independently of SAC signaling. However, the differences in the benomyl sensitivity between two strains expressing *spc105* mutants containing only strong (#1-3) or weak (#4-6) MELT motifs persisted, even when Sgo1 recruitment to the centromere was either decoupled from Bub1 recruitment (*bub1*^Δ^*^kinase^*; Fig. S1E) or completely abolished (*sgo1Δ*; Fig. S1F). It should be noted that the anterior basic patch in Spc105 was also mutated in these strains, because otherwise the strains are inviable on benomyl (Roy et al., 2019). In conclusion, reduced Sgo1 recruitment by the Spc105 mutants does not fully explain the observed differences in their benomyl-sensitivity.

Finally, benomyl affects microtubule dynamics throughout the cell cycle creating the possibility that reasons unrelated to SAC strength contribute to the benomyl sensitivity displayed by the various mutants. To verify that the observed differences among the mutant strains were mainly due to a weaker SAC, we disrupted the mitotic functions of the kinesin Kar3 in strains expressing Spc105 mutants with either strong or weak MELT motifs (Jin et al., 2012). Disruption of the mitotic Kar3 activity impedes chromosome capture by spindle microtubules and also delays bipolar spindle formation (Jin et al., 2012; Tanaka et al., 2005). Consequently, a robust SAC becomes essential for maintaining genome stability. We observed the same correlations between the strength of active MELT motifs in Spc105 mutants and cell viability in this assay as the benomyl sensitivity experiments (Fig. S2A and S2B). Therefore, we conclude that differences in the sensitivity of Spc105 mutant strains mainly arise from differences in their SAC signaling strengths.

### Spc105 mutants containing only weak MELT motifs experience high rates of chromosome missegregation during unperturbed cell division

All the assays so far examine changes in the SAC signaling strength of Spc105 mutants under stressful conditions. To reveal whether the impaired SAC signaling activity due to the presence of only weak MELT motifs in Spc105 translates into chromosome missegregation under normal conditions, we used the colony sectoring assay (Duffy and Hieter, 2018; Hieter et al., 1985). In this assay, chromosome missegregation frequency is quantified for an artificial, linear yeast chromosome. Cells that successfully propagate this chromosome appear white, but turn red/pink in color when the artificial chromosome is lost due to missegregation. Therefore, the frequency of color transformations provides a quantitative read-out of chromosome missegregation events.

We studied the impact of the weak SAC signaling activity displayed by Spc105^#4-6^ (mutant containing motifs #4-6 as the phosphorylatable motifs) under normal growth conditions using the colony color assay. It should be noted that we scored both half-sectored and fully red colonies as chromosome mis-segregation events (see Methods for details). We found that the chromosome missegregation events were significantly more frequent in cells expressing Spc105^#4-6^ compared a wild-type strain. In fact, they were significantly more frequent even when compared to the *mad1*Δ strain lacking a functional SAC (Fig. 2F). This is likely because of impaired chromosome biorientation due to the reduced Sgo1 recruitment in this strain (Vleugel et al., 2015). It is important to note that SAC signaling in this mutant strain is indistinguishable from the SAC signaling in a wild-type strain in the traditional assay using nocodazole treatment, (see Bub3-mCherry recruitment and flow cytometry data in Fig. S3A-B).

These results demonstrate that in cells containing a large number of signaling kinetochores, just one strong MELT motif or several weak MELT motifs per Spc105 can impose a prolonged mitotic arrest. The cumulative signaling output from a large and unchanging number of unattached kinetochores is able to mask significant defects in the signaling outputs of individual kinetochores in this assay. However, to ensure accurate chromosome segregation during normal cell division, Spc105 containing many MELT motifs with a high Bub3-Bub1 binding affinity are essential.

The requirement of many, strong MELT motifs per Spc105 for accurate chromosome segregation raises a critical question: why hasn’t the functionally optimal amino acid sequence retained by all six MELT motifs in Spc105? The explanation is likely to be that many high affinity MELT motifs reduce the responsiveness of the SAC to silencing mechanisms. In budding yeast, the SAC is silenced when the SAC signaling proteins dissociate from the kinetochore, and they are prevented from rebinding by PP1-mediated dephosphorylation. An intriguing hypothesis is that the high affinity MELT motifs are resistant to dephosphorylation, and hence they retard SAC silencing. The structure of the MELpT-Bub3-Bub1^289-359^ complex shows that the phosphorylated threonine residue is lodged in a positively charged groove on Bub3 surface (Primorac et al., 2013). This mode of binding implies that the Bub3-Bub1 complex will sterically interfere with PP1 by shielding the MELT motif from PP1 (see Fig. 1B). Consequently, the rate of dephosphorylation of a MELT motif may be inversely correlated with its Bub3-Bub1 binding affinity, and by extension, its amino acid sequence. Given that only two strong MELT motifs are sufficient for a strong SAC, sequence divergence of some of the MELT motifs may be desirable, because this allows the mutants to trade SAC strength for responsive, achieving timely anaphase onset and hence a rapid growth.

### Kinetochore-microtubule attachment reduces its kinase activity in the kinetochore, priming the SAC for silencing

Before investigating the effects of MELT motif sequence on SAC responsiveness to silencing, we found it necessary to fully understand the mechanisms of SAC silencing itself. In budding yeast, two mechanisms contribute to SAC silencing: (1) the physical separation of the Spc105 phosphodomain from Mps1 kinase bound to the Calponin-Homology domains of the Ndc80 complex (Aravamudhan et al., 2015), and (2) recruitment of the Protein Phosphatase I (PP1) via the conserved ‘RVSF’ motif in N-terminus of Spc105 to dephosphorylate the MELT motifs and disrupt the recruitment of SAC proteins (Liu et al., 2010; London et al., 2012; Meadows et al., 2011b; Rosenberg et al., 2011). We wanted to determine why both mechanisms are necessary for anaphase onset, and more importantly, how their contributions affect the responsiveness of the SAC to silencing.

In budding yeast, Mps1 kinase binds to the CH-domains of Ndc80 even after the formation of end-on attachment, which implies that a gradient of Mps1 kinase activity is likely to exist in bioriented yeast kinetochores (Aravamudhan et al., 2015; Koch et al., 2019). We also found that the native Spc105 phosphodomain extends past its average position toward the high Mps1 kinase activity regions of the kinetochore as evidenced by FRET between donor fluorophore inserted at five locations along Spc105 and strategically placed acceptor fluorophores (Fig. S4A) (Aravamudhan et al., 2014; Joglekar et al., 2009). The persistence of Mps1 in bioriented kinetochores and unstructured nature of Spc105 led us to hypothesize that MELT motifs will continue to be phosphorylated even after biorientation, and this should delay SAC silencing. To test this, we tethered an additional Spc105 phosphodomain at different distances from the CH-domains using rapamycin-induced dimerization of Fkbp12 and Frb in asynchronously growing cells (Fig. 3A). Consistent with our hypothesis, even though the phosphodomain did not activate the SAC when tethered distally from the CH-domains (at the C terminus of either Spc34 or Ndc80), it was robustly phosphorylated as evidenced by the recruitment of Bub3-mCherry (Fig. 3A, also see Fig. S4B). Bub3-mCherry recruitment decreased progressively as the average separation between the tethering point and the CH-domains increased (as measured in bioriented kinetochores, (Joglekar et al., 2009)). The inclusion of the PP1 recruitment motif in Spc105 (+PP1) reduced, but did not eliminate, Bub3 recruitment (+PP1 in Fig. 3A, Fig. S4C). In fact, a sizeable fraction (∼20%) of anaphase cells displayed Bub3-mCherry colocalized with kinetochores. Finally, the phosphodomain recruited Bub3-mCherry even when we tethered it after kinetochore biorientation in metaphase arrested cells indicating that not all of the Bub3 was recruited prior to biorientation (Fig. 3C). Importantly, we found that the phosphodomain did not recruit Mad1 when it was tethered to a position distal to the CH-domains (Fig. 3D).

**Figure 3.**
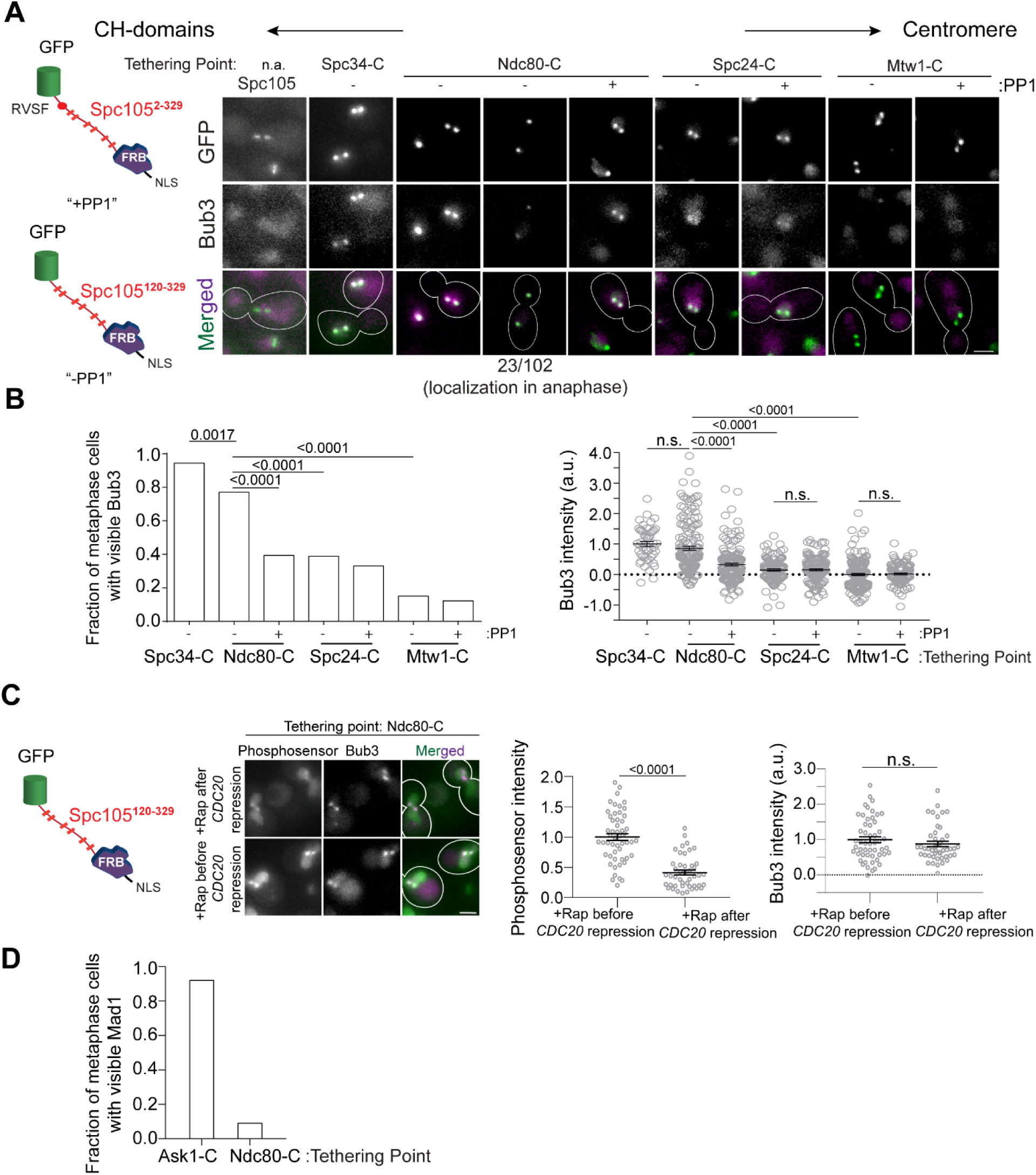
A gradient of Mps1 kinase activity continues to phosphorylate MELT motifs after chromosome biorientation. **A** right: Cartoons show schematics of the two Spc105 phosphodomains tethered to kinetochore subunits using the rapamycin-induced dimerization of Fkbp12 and Frb domains. Micrographs show representative images of cycling cells wherein the phosphodomain is tethered to the C-terminus of the indicated kinetochore subunit. (n.a. - Not Applicable). **B** Left: The bar plot shows the scoring of cells with bioriented kinetochore clusters based on whether or not they visibly recruit Bub3-mCherry. (n=71 for Spc34-C, 109 and 94 for Ndc80-C, 95 and 103 for Spc24-C, 106 and 122 for Mtw1-C, accumulated from two experimental and biological repeats (whenever possible). Right: The scatter plot on the right shows the quantification of Bub3-mCherry fluorescence from all cells with bioriented kinetochore clusters. (mean+s.e.m. n=47 for Spc34-C, 159 and 153 for Ndc80-C, 114 and 154 for Spc24-C, 173 and 104 for Mtw1-C, accumulated from two experimental and biological repeats wherever possible). n.s. Not significant. **C** Micrographs show the recruitment of Bub3-mCherry by the phosphosensor tethered at the Ndc80 C-terminus in cycling (before *CDC20 repression)* and metaphase arrested cells (after *CDC20* repression). Scatterplots show the relative fluorescence signal from the GFP-tagged phosphosensor and Bub3-mCherry respectively. (mean+s.e.m. n=52 and 45 for rapamycin treated samples before and after Cdc20 depletion from two technical replicates). n.s. Not significant. **D** Bar graph shows fraction of cells visibly recruiting Mad1-mCherry as shown in the micrographs (n= 242 and 142 for Ask1-C and Ndc80-C respectively).

Thus, a diminishing gradient of Mps1 kinase activity extending from the CH-domains toward the centromere exists in bioriented yeast kinetochores. This residual Mps1 kinase activity can phosphorylate MELT motifs, but it is likely not sufficient to phosphorylate Bub1, possibly because of PP1 activity. These data suggest that the reduction of Mps1 activity after microtubule attachment primes the SAC for silencing, but this is not sufficient to induce anaphase onset.

### PP1-mediated dephosphorylation of Bub1 contributes to SAC silencing, but this silencing mechanism is not as efficient as PP1-mediated dephosphorylation of MELT motifs

We next examined the role of PP1 in SAC silencing. PP1 can disrupt SAC signaling cascade by dephosphorylating several Mps1 targets in recruitment step of the SAC signaling (Faesen et al., 2017; Ji et al., 2017; London and Biggins, 2014; Tipton et al., 2013) (Fig. 4A). To confirm this, we examined the recruitment of SAC proteins in strains expressing *spc105^RASA^*, an Spc105 mutant that cannot recruit PP1. In cells expressing *spc105^RASA^*, the SAC cannot be silenced, therefore these experiment were performed in strains that lacked a SAC signaling proteins (Rosenberg et al., 2011). As expected, in cells expressing spc105^RASA^ (and carrying suitable SAC gene deletions) Bub1 and Mad1 recruitment persisted even after biorientation (Fig. 4B). In contrast, the tethered phosphodomain recruited Bub1, but not Mad1, suggesting to us that PP1 must dephosphorylate Bub1 to silence the SAC (Fig 3D).

**Figure 4.**
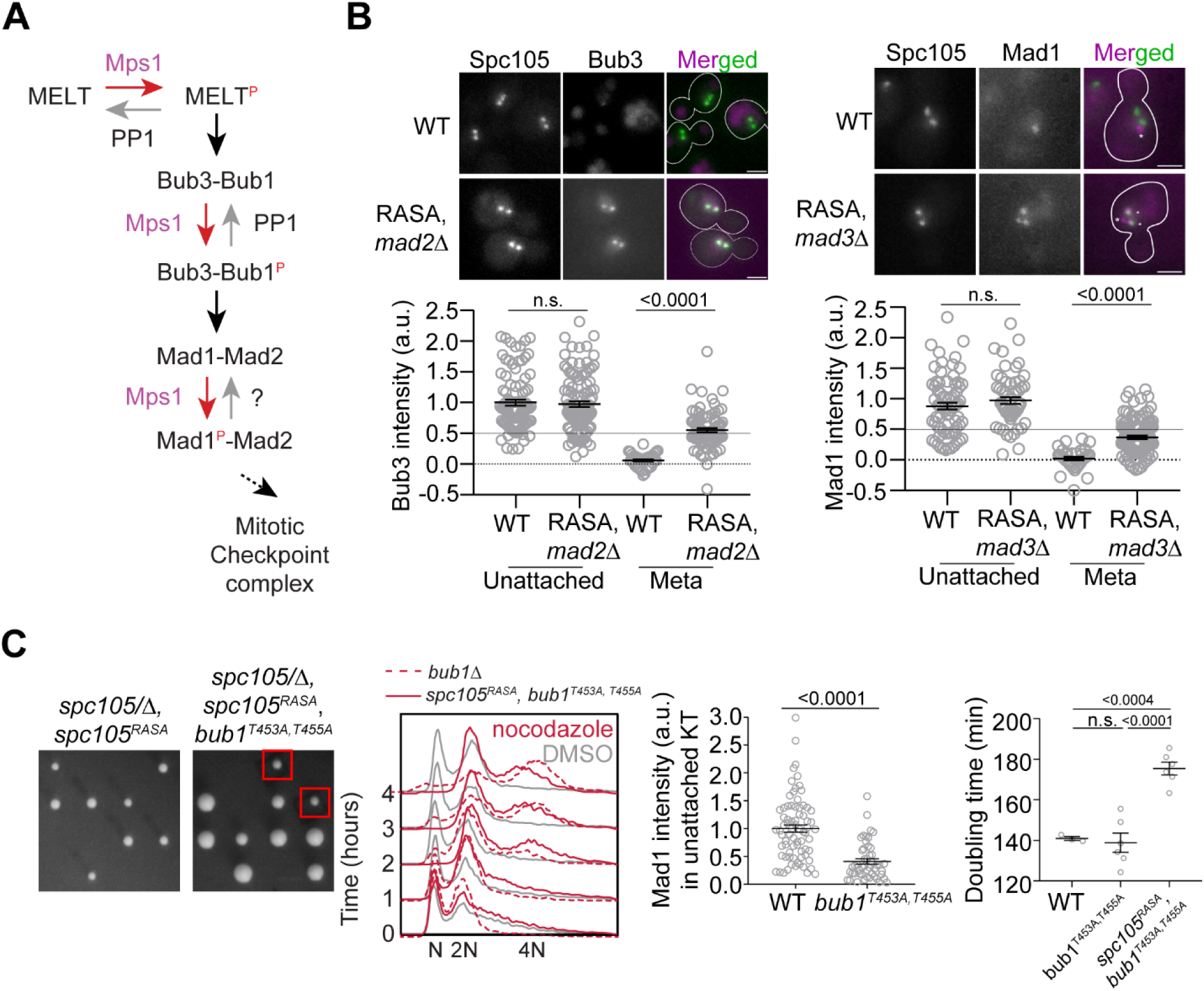
Revealing the determinants of the responsiveness of the SAC to PP1-mediated silencing. **A** Schematic of the SAC signaling cascade highlighting the potential steps that can be disrupted by Protein Phosphatase I (PP1). **B** Top: Representative micrographs of cells with bioriented kinetochore clusters showing the colocalization of the indicated proteins with fluorescently labeled kinetochores. Note that the Mad1-mCherry puncta marked with an asterisk result from the deletion of the nuclear pore protein Nup60. These puncta are not associated with kinetochores. The ones co-localized with kinetochores are marked with arrowheads. Scale bar∼3.2µm. Scatter plots at the bottom show the quantification of fluorescence signal of kinetochore colocalized Bub3-mCherry and Mad1-mCherry (mean+s.e.m., normalized to the respective average signal measured in nocodazole-treated cells). Bub3-mCherry: n=93 and 37 for WT, 107 and 77 for Spc105^RASA^, *mad2*Δ. Mad1-mCherry: n=74 and 34 for WT, 60 and 111 for Spc105^RASA^, *mad3*Δ, pooled from two technical repeats. n.s. Not significant. **C** Left: Tetrad dissection analysis of the indicated strains. Second from the left: Flow cytometry of DNA content of the indicated strains during prolonged exposure to nocodazole. The best of the two technical repeats is shown here. Second from the right: quantitative comparison of Mad1-mCherry recruited by unattached kinetochores in nocodazole-treated cells. (mean+s.e.m. n=74 for WT and 49 for *bub1^453A, 455A^*, pooled from two experimental repeats). Right: Doubling time of the indicated strains in rich media (horizontal line indicates the mean value, obtained from at least three experimental repeats). n.s. Not significant.

To understand the role of PP1-mediated dephosphorylation of Bub1, we created a Bub1 mutant wherein two phosphosites implicated in Mad1 recruitment are replaced with non-phosphorylatable alanine residues, and asked if this mutant rescues the viability of *spc105^RASA^* strain (Ji et al., 2017; London and Biggins, 2014; Qian et al., 2017). The *spc105^RASA^ bub1^453A, 455A^* double mutant was viable, suggesting that the dephosphorylation of Bub1 by PP1 aids SAC silencing (Fig. 4C, plate photographs). Surprisingly, however, the SAC was still active in this double mutant as judged by the ability of this strain to maintain 2N ploidy upon a prolonged exposure to nocodazole (Fig. 4C, 2^nd^ plot from the left). Mad1 recruitment to unattached kinetochores in nocodazole-treated cells was diminished, but not eliminated indicating that other phosphorylation sites in Bub1 also aid in Mad1 recruitment (London and Biggins, 2014) (Fig. 4C, scatter plot, second from the right). The double mutant also grew in benomyl-containing media, which indicates that the *bub1^453A, 455A^* mutation weakens, but does not abrogate, the SAC (Fig. S4D). Importantly, we found that the *spc105^RASA^ bub1^453A, 455A^* double mutant experienced longer doubling times in YPD, which is consistent with a SAC silencing defect arising from the lack of recruitment of PP1 by Spc105 (Fig. 4C, scatter plot on the right). Thus, PP1 must dephosphorylate several phosphorylated residues on Bub1 to completely inhibit Mad1 recruitment.

These experiments clarify the distinct roles of the modulation of Mps1 activity and PP1 and their efficiency in SAC silencing. Attachment-mediated separation of the Spc105 phosphodomain and Ndc80 CH-domains along with partial dissociation of Mps1 from the CH-domains significantly reduces Mps1 kinase activity in the inner kinetochore, and makes SAC signaling cascade responsive to PP1-mediated dephosphorylation. PP1 is then able to dephosphorylate MELT motifs and Bub1 to silence the SAC. However, PP1 must dephosphorylate multiple sites in Bub1 to completely inhibit Mad1 recruitment. In contrast, dephosphorylation of just one residue in the MELT motif prevents the recruitment of both Bub3-Bub1 and Mad1, which implies that the PP1-mediated dephosphorylation of MELT motifs is more efficient than Bub1 dephosphorylation. Consequently, if Bub3-Bub1-binding to the MELT motif interferes with MELT motif dephosphorylation, then the affinity of each MELT motif for Bub3-Bub1, and hence its sequence, will strongly influence the efficiency of its dephosphorylation.

### The number of high affinity MELT motifs in Spc105 and Bub1 expression together determine SAC responsiveness to silencing mechanisms

To reveal the influence of the affinity of MELT motifs on the responsiveness of the SAC to the silencing mechanisms, we created a hyper-active Spc105 variant that we denote as #1^6^. In this mutant, a span of 10 amino acids centered on the last five MELT motif is replaced with the same span centered on the optimal sequence of the first MELT motif. In a large fraction of cells expressing #1^6^, Bub1-mCherry abnormally colocalized with bioriented kinetochore clusters indicating a defect in SAC silencing (Fig. 5A, left and S5A). Accordingly, the doubling time for these strains was ∼ 14% higher compared to wild-type cells (164 vs 144 min.; Fig. 5A, middle scatter plot). The increased doubling time is due to the SAC, because the deletion of *MAD1* eliminated it (5A, second from the right). Importantly, the stronger SAC activity of #1^6^ did not confer any detectable advantage to yeast cells: the growth rate and maximal cell density reached was similar in both wild-type and #1^6^ when the two strains were grown in media containing benomyl (Fig. 5A right). Therefore, the retardation of the cell cycle due to the hyper-active Spc105 allele is a fitness cost for this yeast strain, and too many high affinity MELT motifs per Spc105 are counterproductive.

**Figure 5.**
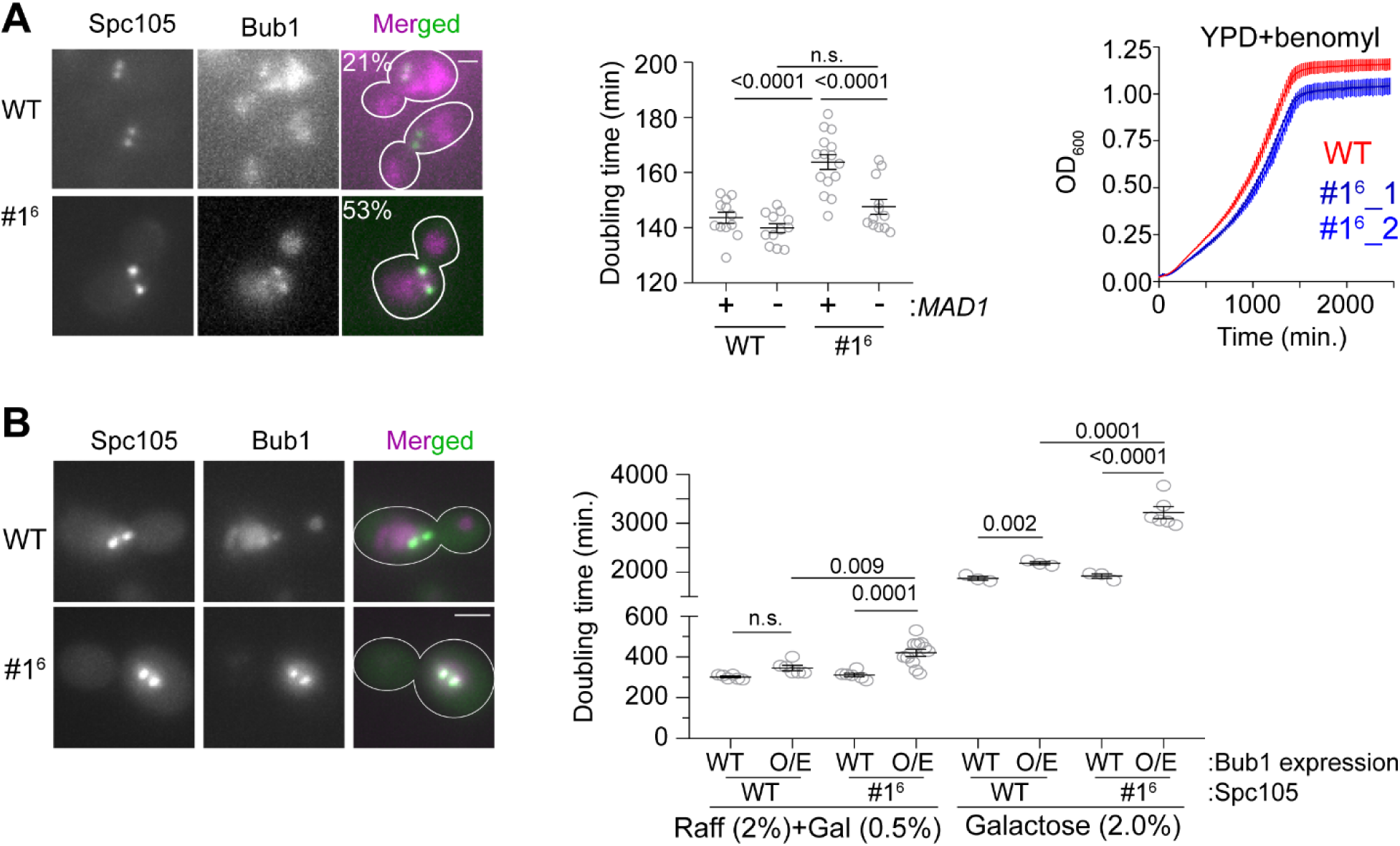
The influence of MELT motif activity and Bub1 expression level on the strength and responsiveness of the SAC. **A** Left: Representative micrographs show the recruitment of Bub1-mCherry to bioriented kinetochore clusters (%age noted at the top of each micrograph indicates the fraction of cells with Bub1-mCherry visibly recruited to bioriented kinetochore clusters; also see Fig. S4A). Scale bar ∼3.2µm. Second from the left: Quantification of the growth rate of indicated strains in non-selective media. Second from the right: The scatter plot shows the indicated strains (mean+s.e.m from at least 6 repeats). Right: Quantification of the growth rate of indicated strains in benomyl-containing media (20µg/ml) (mean+/- s.e.m. from n= 3). n.s. Not significant. **B** Left: Representative images show Bub1 recruitment at the bi-oriented kinetochores when Bub1 is overexpressed in indicated strains (n=2). Scale bar ∼3.2µm. Right: Scatter plot shows the doubling time of the indicated strains, grown in raffinose (2%)+galactose (0.5%) and in galactose (2%). mean+s.e.m obtained from at least 6 repeats. Note that the growth rates of all strains are greatly reduced, when raffinose or galactose rather glucose are used as the carbon source. n.s. Not significant.

The abundance of Bub3-Bub1 is also likely to influence the balance between SAC strength and responsiveness, because Bub1 expression level limits number of phosphorylated MELT motifs occupied by Bub3-Bub1 in unattached kinetochores (Aravamudhan et al., 2016). To test this, we over-expressed Bub1-mCherry using the galactose-inducible *GAL1* promoter, and quantified the rate of growth of cells expressing either wild-type Spc105 or #1^6^ (Janke et al., 2004). Upon partial induction of the *GAL1* promoter using low galactose concentration (0.5%, i.e. moderate over-expression), kinetochores retained Bub1 even after biorientation in both strains (micrographs in Fig. 5B, left). The doubling time of cell cultures of both strains was significantly higher, but the effect was more severe for the #1^6^ culture (Fig. 5B, S5B left). Upon full induction of *GAL1* promoter activity using 2% Galactose, the growth of both strains was significantly hampered (Fig. S5B right and 5B). Moreover, the fold-increase in the doubling time was higher for the #1^6^ cultures compared to wild-type cell cultures in both experiments (∼ 15% for wild-type Spc105 with either partial or full induction of the *GAL1* promoter, 35 and 66% for MELT1^6^ in partial and full induction conditions respectively).

In conclusion, the low expression level of Bub1 limits the recruitment of Bub3-Bub1 and Mad1-Mad2 to unattached yeast kinetochores and thus limits SAC strength. The number and affinity of MELT motifs determines the responsiveness of the SAC to PP1-mediated silencing.

### Engineering an Spc105 allele for optimal SAC signaling, silencing, and error correction

These findings point to a trade-off between implementing a strong SAC and one that is responsive. For a strong SAC, many high affinity MELT motifs are desirable. However, to make the SAC responsive SAC, these MELT motifs must be efficiently dephosphorylated, and this necessitates the recruitment of PP1 by Spc105 via the RVSF motif. This Spc105-PP1 interaction comes with its own cost: the PP1 recruited for SAC silencing can inadvertently stabilize syntelic attachments and thus lead to chromosome missegregation (Roy et al., 2019). Consequently, the weakening or even complete inactivation of the Spc105-PP1 interaction leads to significantly improved chromosome segregation when chromosome biorientation is challenged (Roy et al., 2019). These observations suggested to us a blueprint for the design of an ‘optimal’ Spc105 variant that: (1) avoids the harmful crosstalk between SAC silencing and chromosome biorientation due to the Spc105-PP1 interaction, (2) allows rapid SAC silencing upon chromosome biorientation, and (3) still maintains a strong SAC. To engineer this optimal variant, we anticipated that: (a) the RVSF motif must be inactivated, (b) MELT motifs must have a weak affinity for Bub3-Bub1, and (c) there should be many such MELT motifs per Spc105.

To test our hypothesis, we created two Spc105 mutants that contain a non-functional RVSF motif (*spc105^RASA^*) and either only the first three, strong MELT motifs (designated as #1-3) or only the last three weak ones (designated as #4-6, see Fig. 2). Tetrad dissection analysis of the two diploid strains: *SPC105/spc105*Δ *leu2/leu2*∷*spc105^RASA,#1-3^* and *SPC105/spc105*Δ *leu2/leu2 ∷ pc105^RASA,#4-6^* revealed that only the weaker MELT motifs suppressed the lethality caused by the non-functional RVSF motif (Fig. 6A, also see Fig. S5C). Thus, diffusive interactions between PP1 and Spc105 are sufficient to dephosphorylate the weak MELT motifs, but not for the strong motifs. The simplest explanation this observation is that Bub3-Bub1 binding to the MELT motif hampers the ability of PP1 to dephosphorylate the MELT motif.

**Figure 6.**
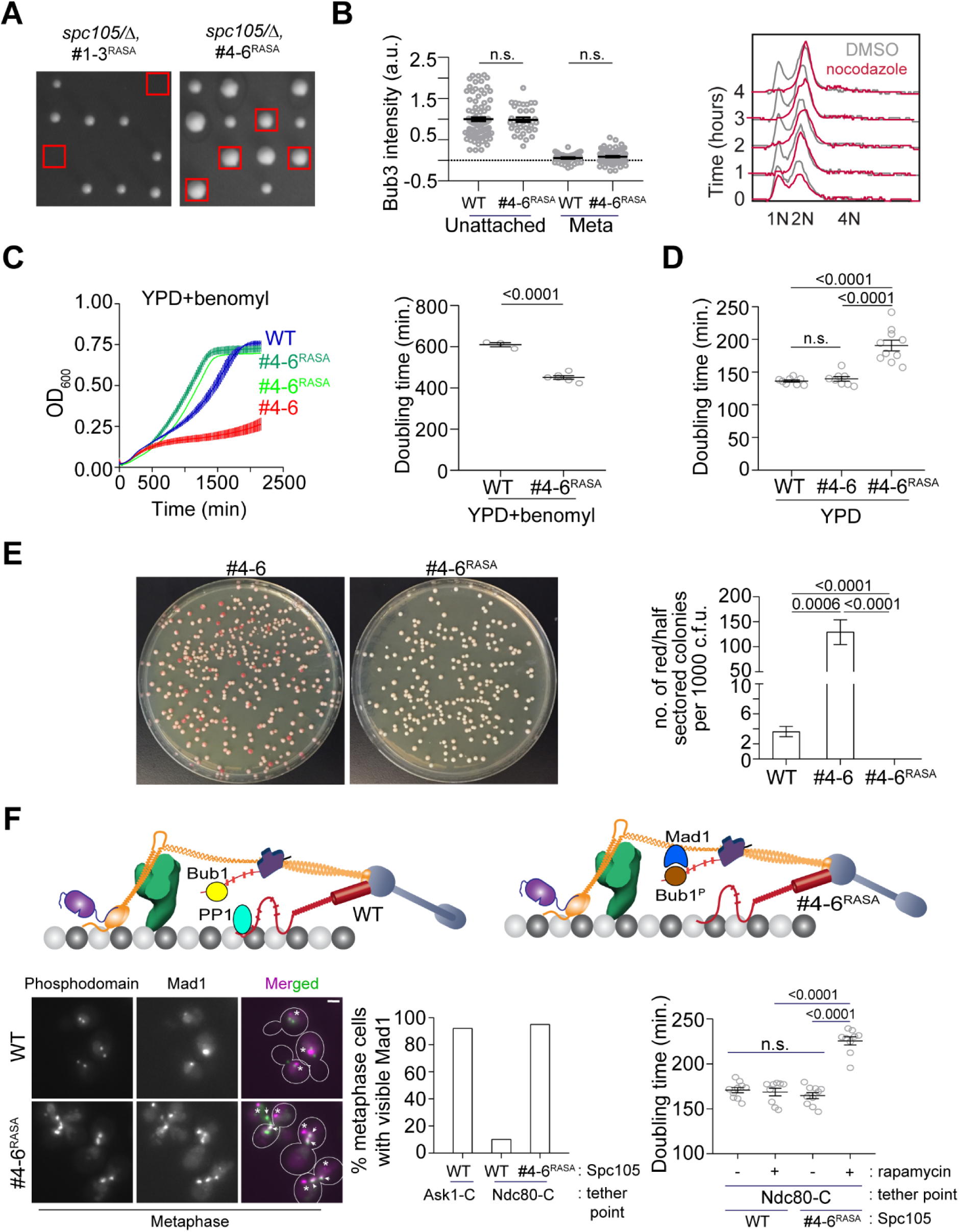
Designing an Spc105 variant optimized for SAC signaling, error correction, and SAC silencing. **A** Left: Tetrad dissections showing the rescue of Spc105^RASA^ by inactivating the first three MELT repeats. Right: Scatter plot depicting Bub3 intensities in unattached and bioriented kinetochores of indicated strains (mean+s.e.m. n=92 and 37 for WT, 33 and 98 for #4-6^RASA^, pooled from two technical repeats). n.s. Not significant. **B** Representative flow cytometry-based quantification of the DNA content of the indicated strains during a prolonged exposure to nocodazole (n=2). **C** Quantification of the doubling times of the indicated strain in YPD respectively (horizontal line marks the mean value, minimum number of experimental repeats n=2). **D** Left: Quantification of population growth for the indicated strains in presence of benomyl (30µg/ml) (mean± s.e.m., n=3). Right: Quantification of the doubling times of the indicated strain in YPD+benomyl (30µg/ml) respectively (horizontal line marks the mean value, minimum number of experimental repeats n=2). **E** Estimation of chromosome loss by colony sectoring assay. Left: Plate images of #4-6 and #4-6^RASA^ strains grown in YPD plates. Right: Bar graph displays the number of red or half sectored colonies per 1000 colony forming units. N= 5081, 4965 and 6596 for WT, #4-6 and #4-6^RASA^ respectively, pooled from ≥ 6 technical repeats. **F** Top: Model explaining why Spc105 phosphodomain tethered in the inner kinetochore does not activate the SAC. Bottom left: Representative micrographs display Mad1-mCherry localization relative to bioriented kinetochores in the indicated strains after treatment with rapamycin. The Mad1-mCherry puncta marked with an asterisk result from the deletion of the nuclear pore protein Nup60. These puncta are not associated with kinetochores. The ones co-localized with kinetochores are marked with arrowheads. Scale bar∼3.2µM. Bottom middle: Bar graph of the percentage of cells visibly recruiting Mad1-mCherry (n= 242, 142 and 290 for Ask1-C, Ndc80-C and Ndc80-C, #4-6^RASA^ respectively, derived from two technical repeats). Bottom right: Scatter plot presents the doubling time of the indicated strains. mean+s.e.m from ≥ three experimental repeats. n.s. Not significant.

Using fluorescence microscopy, we confirmed that kinetochores in the *spc105^#4-6^* strain did not recruit Bub3-mCherry in metaphase, but recruited both Bub3 and Mad1 when unattached upon nocodazole treatment (Fig. 6B, also see Fig S5D and S5E). As expected, the SAC was active under the same condition as evidenced by the quantification of the cellular DNA content (Fig 6B). In benomyl-containing media, this strain grew significantly faster even compared to wild-type cells, suggesting that chromosome biorientation and chromosome segregation accuracy was improved (6D). Colony sectoring assay revealed that even under normal growth conditions, the chromosome segregation accuracy in this strain was significantly higher than the strain expressing *spc105^#4-6^* and even wild-type cells (only 3 red colony out of ∼ 6600 compared to 18 red colonies out of ∼5100 for the wild-type strain). However, the doubling time for this mutant under normal conditions was 20% higher than wild-type cells, (Fig. 6C and S5F). The slowing down of the cell cycle in this case consistent with an SAC silencing defect, but it is accompanied by improved chromosome segregation. These observations demonstrate that the balance of SAC signaling strength and its responsiveness to silencing mechanisms determines the trade-off between accurate chromosome segregation and timely anaphase onset.

### PP1-mediated dephosphorylation of Bub1 prevents the tethered Spc105 phosphodomain from activating the SAC from inner kinetochore

Construction of the *spc105^RASA,#4-6^* strain also afforded us the opportunity to determine why the SAC is not activated, when the Spc105 phosphodomain is tethered in the inner kinetochore (e.g. to Ndc80 C-terminus) even though it robustly recruits Bub3-Bub1 (Fig. 3). We reasoned that the PP1 recruited by Spc105 dephosphorylates Bub1 bound to the tethered phosphodomain preventing it from recruiting Mad1 and thus ensuring that the SAC remains inactive (cartoon in Fig. 6E, top). To confirm this, we tethered the Spc105 phosphodomain to Ndc80-Fkbp12 in *spc105^RASA, #4-6^* cells by adding rapamycin to growth media, reasoning that the Bub1 recruited by the tethered phosphodomain would now remain phosphorylated in the absence of PP1 and thus recruit Mad1. Consistent with this expectation, we found that the phosphodomain tethered at Ndc80-C robustly recruited Mad1 in cells expressing *spc105^RASA, #4-6^* but not wild-type Spc105 (Fig. 6E, representative images at the middle, bar graph at the bottom left). We even observed a small number of cells with wherein Mad1 localized at the kinetochores even in anaphase (S5G). The growth rate of cells expressing the *spc105^RASA, #4-6^*, but not wild-type Spc105 in media containing rapamycin was also reduced, revealing incomplete SAC silencing (6E, bottom right). These experiments demonstrate that the reduction of Mps1 kinase activity in the kinetochore primes the SAC for silencing, enabling PP1 to disrupt SAC protein recruitment.

## Discussion

This study provides three critical insights into the physiological operation of the SAC signaling cascade. First, we show that reduced SAC strength stemming from Spc105 containing fewer or only weak MELT motifs is apparent only in normally growing cells or in cells wherein the process of chromosome biorientation is challenged. It is not detected in nocodazole-treated cells containing a large and constant number of unattached chromosomes. The most likely explanation for this discrepancy is that the cumulative signaling output of the large and unchanging number of unattached kinetochores in nocodazole-treated cells masks the reduced output of individual kinetochores. Therefore, SAC strength determined by measuring the duration of mitotic arrest in nocodazole-treated cells is not useful in predicting the ability of the SAC to ensure genome stability. Under normal conditions, the maintenance of genomic stability in budding yeast requires a strong SAC enabled by many MELT motifs possessing the functionally optimal amino acid sequence. This insight provides a functional explanation for the evolutionary trend of the expansion of the number of MELT motifs in Spc105 and its homologs via recurrent duplication events (Tromer et al., 2015).

Second, our study clarifies the distinct roles that the microtubule-mediated reduction in Mps1 activity and PP1 play in silencing the SAC. The microtubule-mediated separation and dissociation of Mps1 significantly reduces its kinase activity, especially in the inner kinetochore, beyond the Dam1 complex (Aravamudhan et al., 2015; Hiruma et al., 2015; Ji et al., 2015). This primes the SAC for silencing but doesn’t completely stop it. The residual Mps1 activity can drive SAC signaling at a lower level, which retards mitotic progression. Reduction in Mps1 activity enables the PP1 recruited by Spc105 to dephosphorylate MELT motifs and Bub1, which stops the recruitment of Bub1 and Mad1 to the kinetochore. Although Bub1 dephosphorylation contributes to SAC silencing, MELT motif dephosphorylation by PP1 has a stronger impact on SAC silencing, consistent with the foundational role of the MELT motifs in the SAC signaling cascade. With this insight, we demonstrate that the number and activity of MELT motifs can be suitably balanced to make the dedicated mechanism for PP1 recruitment via Spc105 non-essential. In fact, the elimination of this interaction has a significant advantage: chromosome biorientation and error correction are improved under stressed conditions, however this comes at the cost of delayed SAC silencing under normal conditions.

The final insight from our work is that the Bub3-Bub1 binding affinity of MELT motifs is a key determinant of the responsiveness of the SAC. This is likely because when the MELT motif is bound by Bub3-Bub1, steric effects interfere with its dephosphorylation. Consequently, the high affinity MELT motifs are more resistant to PP1-mediated dephosphorylation, and they reduce the responsiveness of the SAC to silencing. It is important to note that at a minimum, the signaling output of one unattached kinetochore has to be sufficient only to delay anaphase onset. Beyond this limit, further increase in its signaling capacity does not offer additional advantage. Therefore, beyond a certain number, additional MELT motifs per Spc105 are unlikely contribute to the strength of the SAC. Instead, they will unnecessarily delay anaphase onset, and thus impose a fitness cost. Thus, the number and Bub3-Bub1 binding affinity of MELT motifs must be optimized to balance SAC strength and responsiveness. Therefore, we propose that the evolution of MELT motif copy number and sequence may reflect an evolutionary process that balances SAC strength and responsiveness to ensure genomic stability and timely cell division.

## Acknowledgements

This work was funded by the GM095005 from NIGMS. We thank Prof. Zuzana Storchova (Technical University Kaiserslautern, Germany) for sharing reagents and Prof. Mara Duncan and her lab (Department of Cell and Developmental Biology, University of Michigan Medical School) for help with the plate reader assay. We also thank Iain Cheeseman, Mara Duncan, and Yukiko Yamashita for providing feedback on this study and the manuscript.

## Materials and methods

### Plasmid and strain construction

The plasmids and *S. cerevisiae* strains used in this study are listed in supporting tables S1 and S2 respectively. Strains containing multiple genetic modifications were constructed using standard yeast genetics. Proteins tagged with GFP(S65T) and mCherry or yeast codon optimized mCherry were used to visualize kinetochores, spindle pole bodies and SAC signaling components. A 7-amino-acid peptide (sequence: ‘RIPGLIN’) was used as the linker between the proteins and their C-terminal tags (GFP, mCherry or 2XFKBP12). The cassettes for gene deletion, gene replacement and C-terminal tags were introduced at the endogenous locus through homologous recombination of PCR amplicons or using linearized plasmids (Jürg Bähler et al., 1998). We previously observed a significant strain to strain variation in the intensity of mCherry-tagged kinetochore proteins or checkpoint proteins due to inherent variability of the brightness of mCherry. Therefore, we created all Bub3-mCherry and Mad1-mCherry strains by crossing the same transformant of either Bub3-mCherry (AJY3844 or AJY3846) or Mad1-mCherry (AJY1836 or AJY3741) with other strains. For the same reason, we created all the strains used in Forster Resonance Energy Transfer (FRET) quantification by crossing a specific transformant of Spc25-mCherry (AJY778) or Dad4-mCherry (AJY906). The deletion mutant of *NUP60* always accompanies Mad1-mCherry to disrupt Mad1localization to the nuclear envelopes (Scott et al., 2005). This facilitated clearer visualization and quantification of Mad1 localized to unattached kinetochores without affecting SAC strength.

To construct diploid strains, we mixed overnight grown cultures of two strains of A and α mating types and spotted the cell suspension on a YPD plate, and then incubated the cells for approximately 3-4 hours at 32°C. To induce meiosis, we transferred stationary phase diploid cells to starvation media (yeast extract 0.1%, Potassium acetate 1%), and we incubated them at RT for 4-5 days to induce meiosis.

All the Spc105 mutants used in the study are chimeras of Spc105 and either GFP or codon optimized mCherry. Genes encoding the chimeric proteins were introduced using a cassette that consists of the 397 bp upstream and 250 bp downstream sequences of the SPC105 open reading frame as promoter (*prSPC105*) and terminator (*trSPC105*) sequences respectively. We introduced genes encoding either GFP (S65T) or a codon optimized mCherry at the 222^nd^, 455^th^, and 709^th^ amino acid position of Spc105 by sub-cloning where we introduced an extra *Bam*HI site (Gly-Ser) at the upstream and *Nhe*I site (Ala-Ser) at the downstream of the GFP fragment. The plasmids based on pRS305 or pRS306 backbone were linearized by *Bst*EII or *Stu*I before transformations to ensure their integration at the *LEU2* or the *URA3* locus respectively.

We constructed the MELT motif mutants of Spc105 using the QuickChange II XL Site-Directed mutagenesis kit (Agilent Technologies). To build pAJ737 (Spc105_#4-6 in #1-3), we cloned a 306 bp fragment which codes for Spc105^223-324^ within *Bam*HI-*Bsi*WI sites of pAJ613 (Spc105^#1-3^). To generate pAJ747 (Spc105^#1-3 in #4-6^), we used pAJ609 (Spc105_#4-6) as vector where we cloned a 274 bp fragment within *Nhe*I-*Mlu*I, that codes for Spc105^132-222^. While constructing the chimeras for #2^MELT^ (pAJ867), #4^MELT^ (pAJ868) and #5^MELT^ (pAJ870), we introduced mutations of M171L, ID232-233ME and I283L respectively, but kept the flanking sequences unchanged. To design a plasmid that codes for Spc105_#1^6^ (pAJ904), we used pAJ419 (Spc105^455::GFP^) as the parent vector. We used *Bsi*WI-*Bam*HI sites in that plasmid to replace the WT sequence which contains 6 MELT repeats with a 981 bp fragment which harbors 6 repeats of MELT1 and its flanking sequences (codes for PDTSGMELTEPIVATP).

To construct pAJ852 or pAJ896, which express *bub1^T453A, T455A^*, we used the pSK954 plasmid backbone (Kemmler et al., 2009). pSK954 consists of the *ADH1* transcription terminator cloned within *Asc*I-*Bgl*II. We cloned the 500bp upstream sequence that harbors *BUB1* promoter, 3.063kb *BUB1* ORF sequence and 651bp *2XFKBP12* within *Sac*II-*Asc*I site of this plasmid. The ORF and *2XFKBP12* are separated by 21bp linker which codes for RIPGILK. We also cloned 350bp downstream sequence of *BUB1* which consists of *BUB1* terminator within *Pme*I-*Apa*I. To build the strains with *bub1^T453A, T455A^* allele, we first created a diploid strain where one copy of *BUB1* was deleted with *TRP1*. The plasmids of pAJ852 or pAJ896 were digested by *Apa*I and SacI to release 6.279kb fragment which recombined at the deleted bub1 locus replacing the *TRP1* cassette. There is mutation of SR (449-450)TG in *BUB1* ORF of pAJ852. However, upon testing there were no discernable phenotypic differences between the strains constructed by pAJ852 and pAJ896.

We obtained pAJ669 (BZ427) from Prof. Storchova’s lab (Peplowska et al., 2014), which we digested by *Stu*I and transformed in yeast to obtain strains required for CIK1-CC overexpression.

### Cell culture

Yeast strains were grown in YPD (yeast extract 1%, peptone 2%, dextrose 2%) or YPRG (yeast extract 1%, peptone 2%, Raffinose 2%, Galactose 0.5%) at 32°C. Strains that express *CDC20* from *MET3* promoter were grown in synthetic dextrose media lacking methionine were used.

For the spotting assays involving CIK1-CC overexpression, we grew all the strains to mid-log phase in YPR media (yeast extract 1%, peptone 2%, Raffinose 2%). Then starting from 0.2 OD_600_ cells, we prepared serial dilutions (1:2 or 1:10) which were then spotted on either YPD or YPG agar media. All spotting assays were performed with at least two biological replicates whenever possible and two technical replicates on both YPD and YPG plates. For time course imaging experiments involving CIK1-CC overexpression, we started from overnight inoculums grown in YPR, shifted the cells to YPG media for 1.5h, and then supplemented the media with α factor (2μg/ml) to arrest the cells at G1 for 2h. After this, we washed the cells to remove the α factor and released them in fresh YPG media. We took aliquots of cells to image and analyze 75, 105 and 135mins after the wash. For chromosome loss assay, the strains containing the nonessential linear chromosome fragment (CFIII, which contains *SUP11*) were grown in synthetic media devoid of uracil until they were diluted for plating.

### Benomyl sensitivity assay

The assay was done as described previously (Roy et al., 2019). We used plates with both 20μg/ml and 30μg/ml benomyl concentration. At least two biological replicates wherever possible and two technical replicates were conducted in all the experiments. We included a wild-type positive control which grows on benomyl, and an appropriate negative control strain which either grows poorly or not at all under the same condition. We prepared benomyl supplemented YPD as it was described previously (Gupta et al., 2018). To track biorientation and spindle formation in benomyl-containing media, we arrested cells in G1 by exposing mid-log phase cultures to α factor (2 μg/ml) for 1h 45min. G1 synchronized cells were then washed to remove the α factor, and transferred to media containing benomyl. Aliquots of these cultures drawn at 45, 75, and 105min post-wash were used for imaging. At least 20 microscope fields were captured at each time point for analysis.

### 96 well plate liquid culture assay

This assay was conducted as described previously (Hung et al., 2018). Briefly, we initiated cultures at 0.05 OD_600_ in each well by appropriately diluting mid-log phase cultures maintaining ∼ 160 μl final volume. For assay involving benomyl treatment, cells from mid-log phase cultures were pelleted, resuspended and diluted in YPD+benomyl liquid. For each strain, we set at least three 3 technical repeats in YPD or YPD+benomyl. In assays that involve rapamycin treatment, we grew the strains to mid log phase in non-selective media, and then diluted these cultures appropriately so as to initiate 0.05 OD_600_ cultures in each well grow in either YPD or YPD+rapamycin (1µg/ml). To measure OD_600_ continuously, we placed the 96 well plate in a Spectra Max 340PC plate reader and incubated it for either 24h (YPD) or 36h (YPD+benomyl) at 30°C without shaking. The reader measured the absorbance every 20mins. For cells growing in YPRG or YPG, we set at least three technical replicates and ran the assay for 48h at 30°C without shaking. It should be noted that the growth rate of these static cultures in a 96 well plate is significantly slower than that of the same cultures grown in a flask, incubated in 30°C shaker incubator.

### Microscopy and image acquisition

A Nikon Ti-E inverted microscope with a 1.4 NA, 100X, oil-immersion objective was used for all the experiments. The 1.5X opto-var lens to measure Bub1, Bub3 and Mad1-mCherry intensities. The cells were imaged at room temperature in synthetic dextrose (or synthetic galactose media whenever it was required for the assay) supplemented with essential amino acids to obtain at least 20 microscopic fields at a given time points for any strains. We added nocodazole or methionine to the mounting media to image the nocodazole arrested cells or Cdc20 depleted cells respectively. For each field of view, a ten-plane Z-stack was acquired (200nm separation between adjacent planes), and at least 20 fields were acquired in each experiment.

Total fluorescence intensities of kinetochore clusters (16 kinetochores in metaphase) was measured by integrating the intensities over a 6×6 region centered on the maximum intensity pixel. We utilized the median intensity of pixels immediately surrounding or a nearby 6×6 area to correct for background fluorescence. The calculations were performed using ImageJ or semi-automated MATLAB programs as described earlier (Joglekar et al., 2006). Microscopy, image acquisition and FRET quantification were performed as described previously (Aravamudhan et al., 2014; Joglekar et al., 2013).

### Chromosome loss quantification

We performed the colony sectoring assay as described previously (Duffy and Hieter, 2018; Warren et al., 2002). We grew strains harboring the nonessential linear chromosome fragment (CFIII, which contains *SUP11*) to mid-log phase on selective media (synthetic media devoid of Uracil), and then plated ∼200-300 cells per plate on non-selective media (YPD). The low adenine concentration in YPD aids the development of red pigments only in cells that lose the CFIII fragment; cells that propagate CFIII remain white in color. We incubated the plates at 30°C for 3 days followed by incubation at 4°C for 24 h which augment the accumulation of the red pigments. To quantify the chromosome loss rate accurately, only those colonies that are either completely red or at least half red/half-sectored, i.e. one half of the colony is red while the other half is white, are counted, because they reflect a chromosome missegregation event in the first cell division immediately after plating. However, because the artificial chromosome is lost only 1 in 10,000 cell divisions (Duffy and Hieter, 2018), a very large number of cells must be plated. We expected our strain expressing the engineered *SPC105* allele to have an even smaller chromosome loss rate than wild-type strains. Therefore, we counted colonies that are either fully red or half-sectored. A fraction of the red colonies represents cells that had already lost the chromosome during the 2-3 cell divisions that occurred during growth to mid-log phase in media selecting for the CFIII fragment. Therefore, the chromosome loss rate reported in this study is expected to be significantly higher than the true chromosome loss rate of the wild type cells that maintain CFIII fragment.

Plate images were acquired using an iPhone 6 camera (Model # MG632LL/A, https://support.apple.com/kb/SP705?locale=en_US). The total number of colonies and the number of red colonies were counted using a custom application (app) written in Matlab. In this app, the total number of colonies on a plate was determined by intensity-based thresholding (after median and top-hat filtering to remove background variation) followed by feature segmentation using the Watershed algorithm to separate overlapping colonies or colonies that touch one another. To count the number of red colonies, the plate image was first transformed to the L*a*b color space and then thresholded along the red-green color axis. The code for this app will be made available upon request.

### Statistical methods

To prepare the scatter plots of Bub1, Bub3 or Mad1 intensities, we normalized the data with the mean intensities obtained for wild-type controls in each experiment. We imaged each strain at least twice to obtain significant number of metaphase cells (>50) to quantify the frequency of metaphase cells with visibly recruited SAC proteins (Bub1, Bub3, or Mad1) at the kinetochores in a cell population. We set the scoring analysis so that it divides the metaphase cell population into two groups: cells with visible SAC protein localization and the cells without detectable localization. We did not notice any significant variation in the fraction of metaphase cells of a specific strain with visible SAC protein localization from individual experiments. Hence, we pooled observations from all the experiments and applied Fisher’s exact test in Graphpad Prism (version 8) to determine the statistical significance of these scoring data. The number of cells analyzed for each strain and number of experimental replications is noted in the figure legends. We applied either t-test or one-way anova or two-way ANOVA test to ascertain the statistical significance of the rest of the data using Graphpad Prism (version 8). The p-values obtained from these tests are indicated in the figures. We performed linear regression analyses on the growth curves to obtain the slope of the linear section of each plot of each strains in non-selective media. The doubling time was calculated as: Doubling time= [Log_10_(2)]/slope.

### Flow cytometry

For these experiments, we started from overnight inoculum the designated strains to obtain mid log phase cultures. We added Nocodazole to the media (final concentration 15μg/ml) to depolymerize the spindle and activate the SAC) and rapamycin (1 µg/ml) to induce the dimerization of FRB and FKBP fused proteins (Gillett et al., 2004). We collected samples containing approximately 0.1 OD_600_ cells at 0, 1, 2 and 3 h post drug addition, fixed the cells using 70% ethanol, and stored them in 4°C overnight. Next day, we washed out the ethanol, and treated the samples with bovine pancreatic RNase (Millipore Sigma, final concentration 170ng/µl) at 37°C for at least 6h in RNase buffer (10mM Tris pH8.0, 15mM NaCl). Then we removed the RNase and resuspended the cells in phosphate buffered saline (PBS) and stored them in 4°C. These samples were incubated in Propidium Iodide (Millipore Sigma, final concentration 5µg/ml in PBS) for 1h at RT on the day of the assay. The stained cells were then analyzed using the LSRFortessa (BD Biosciences) in Biomedical research core facility, University of Michigan medical school. We repeated flow cytometry for each strain at least twice. The data was analyzed using the FlowJO software.

---

**Figure S1 (related to Figure 2).**
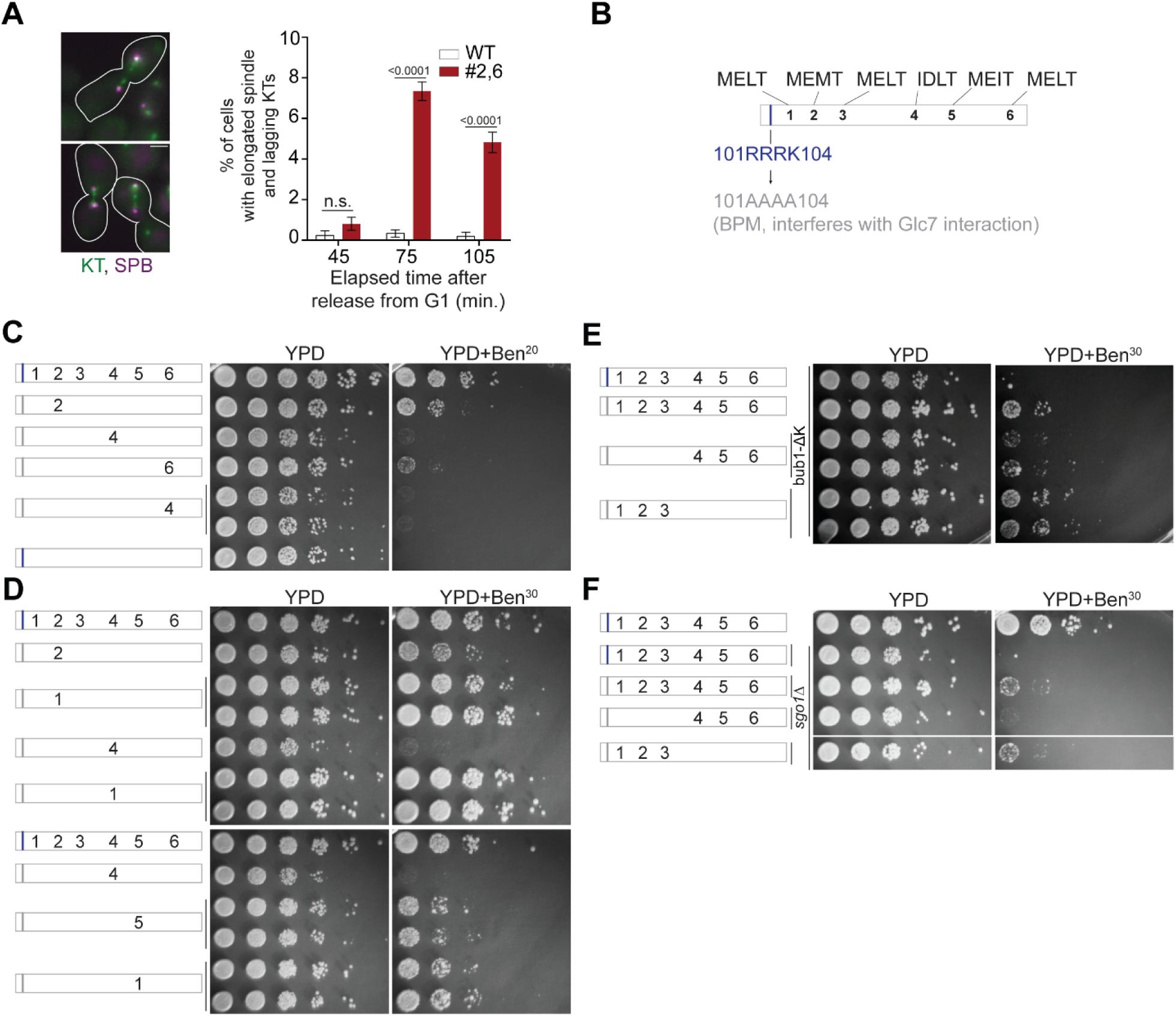
Requirements of the MELT motif number per Spc105 and their affinity for survival on media containing benomyl. **A** Quantification of cells of indicated strains showing an elongated spindle, which is indicative of anaphase onset. (mean+s.e.m., n=1029, 889 and 992 at 45, 75 and 105 min respectively for WT, n=890, 864 and 812 at 45, 75 and 105 min respectively for #2,6, pooled from three technical repeats). n.s. Not significant. **B** Schematic of the Spc105 phosphodomain with the amino acid sequence indicated at the top. The 101-RRRK-104 span is designated as the anterior basic patch of Spc105. It is implicated in the binding of PP1 to Spc105 (Roy et al., 2019). **C-F** Benomyl-sensitivity of the indicated strains. Schematic on the left displays the number and position of active MELT motifs. The gray bar indicates that the basic patch is replaced by non-polar alanine residues (101-RRRK-104::AAAA). **C** We previously found that mutants expressing Spc105-5A are inviable in presence of benomyl (Aravamudhan et al., 2016). Mutation of the basic patch (gray bar) suppresses that sensitivity of Spc105-5A variants only if the MELT motif possesses the optimal, consensus sequence. The benomyl sensitivity of the sixth MELT motif in this assay is surprising given that this motif possesses the consensus amino acid sequence. We speculate that the lower activity is due to the absence of the negatively charged residue directly downstream from the MELT motif, which contributes to the interaction of the MELT motif with the Bub3-Bub1 complex (Primorac et al., 2013). **D** The activity of a MELT motif is determined by its sequence, but not position. **E-F**, The benomyl sensitivity of the #1-3 vs #4-6 in the background of basic patch mutation (101-RRRK-104::AAAA) with either *bub1*^Δ^*^kinase^* or *sgo1*Δ. The basic patch mutation is necessary, because it alleviates the growth of bub1-Δ and *sgo1Δ* in benomyl containing media (Roy et al., 2019).

**Figure S2 (related to figure 2).**
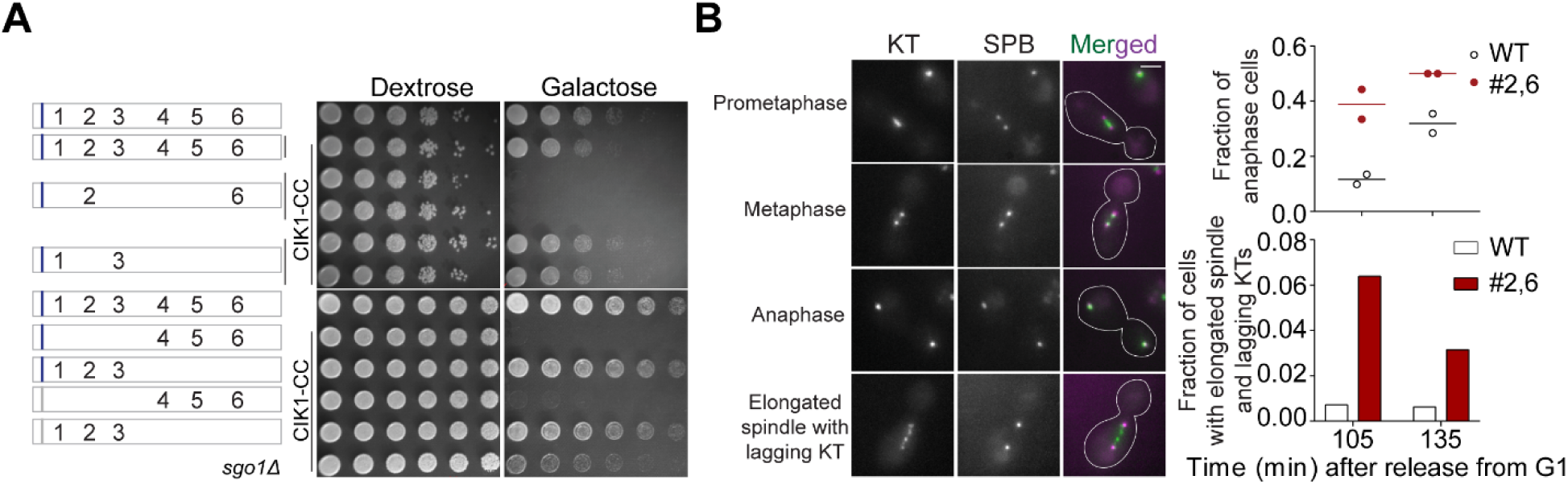
Multiple, strong MELT motifs per Spc105 are essential for cell survival when chromosome biorientation is challenged by conditionally disrupting the mitotic Cik1 functions. **A** Effect of Cik1-CC overexpression (stimulated by the use of galactose as the carbon source) on the indicated strains. **B** Micrographs on the left show the scoring scheme used to identify cells in anaphase and those with elongated spindles, which are indicative of anaphase onset (but cannot be confirmed as anaphase). Bar∼3.2µm. Scatter and bar plots on the right show quantification. (n=694 and 476 at 105 and 135 min respectively for WT and 547 and 317 at 105 and 135 min respectively for #2,6 from 2 repeats).

**Figure S3 (related to Figure 2).**
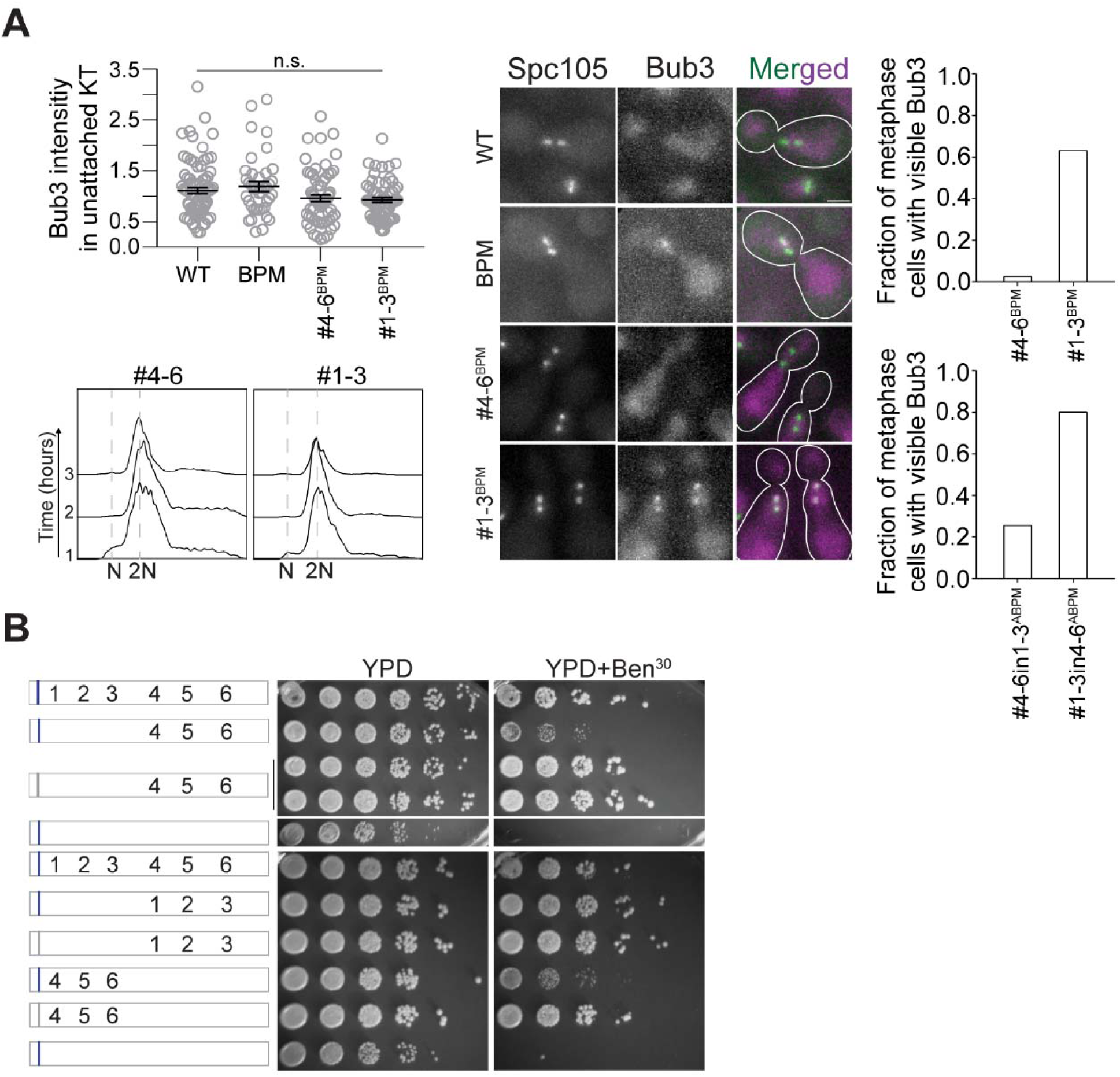
The difference in the SAC signaling activities of the first three and last three MELT motifs in Spc105 is apparent only on benomyl-containing media. **A** Top left: Scatter plot revealing the intensity of Bub3-mCherry in unattached kinetochores of indicated strains when the cells were treated with nocodazole for 2h. (mean+s.e.m. n=82, 39, 67 and 59 for WT, BPM, #4-6^BPM^ and #1-3^BPM^ respectively, accumulated from two repeats). Bottom left: Representative flow cytometry-based quantification of the DNA content of the indicated strains during a prolonged exposure to nocodazole. Middle: Representative images show the recruitment of Bub3-mCherry to bioriented kinetochore clusters in the indicated strains. Bar∼3.2µm. Right: Bar graphs show the fraction of metaphase cells with Bub3 recruited at the kinetochores in *CDC20* depleted (top) and cycling (bottom) population of the indicated strains. Top: n=116 and 106 for #4-6^BPM^ and #1-3^BPM^ respectively, pooled from two biological replicates for each strain. Bottom: n=90 and 96 for #4-6in1-3 and #1-3in4-6 respectively, accumulated from two biological replicates for each strain. n.s. Not significant. **B** Benomyl spotting assay of indicated strains. Schematic on the left shows the number and position of active MELT motifs. The basic patch at the N-terminus is indicated by the blue bar; gray bar indicates that the basic residues in this patch are replaced by non-polar alanine residues. Mutation of anterior basic patch alleviates the growth of strains expressing Spc105^#4-6^, independently of the positions of #4-6 in Spc105 phosphodomain.

**Figure S4 (related to Figure 3).**
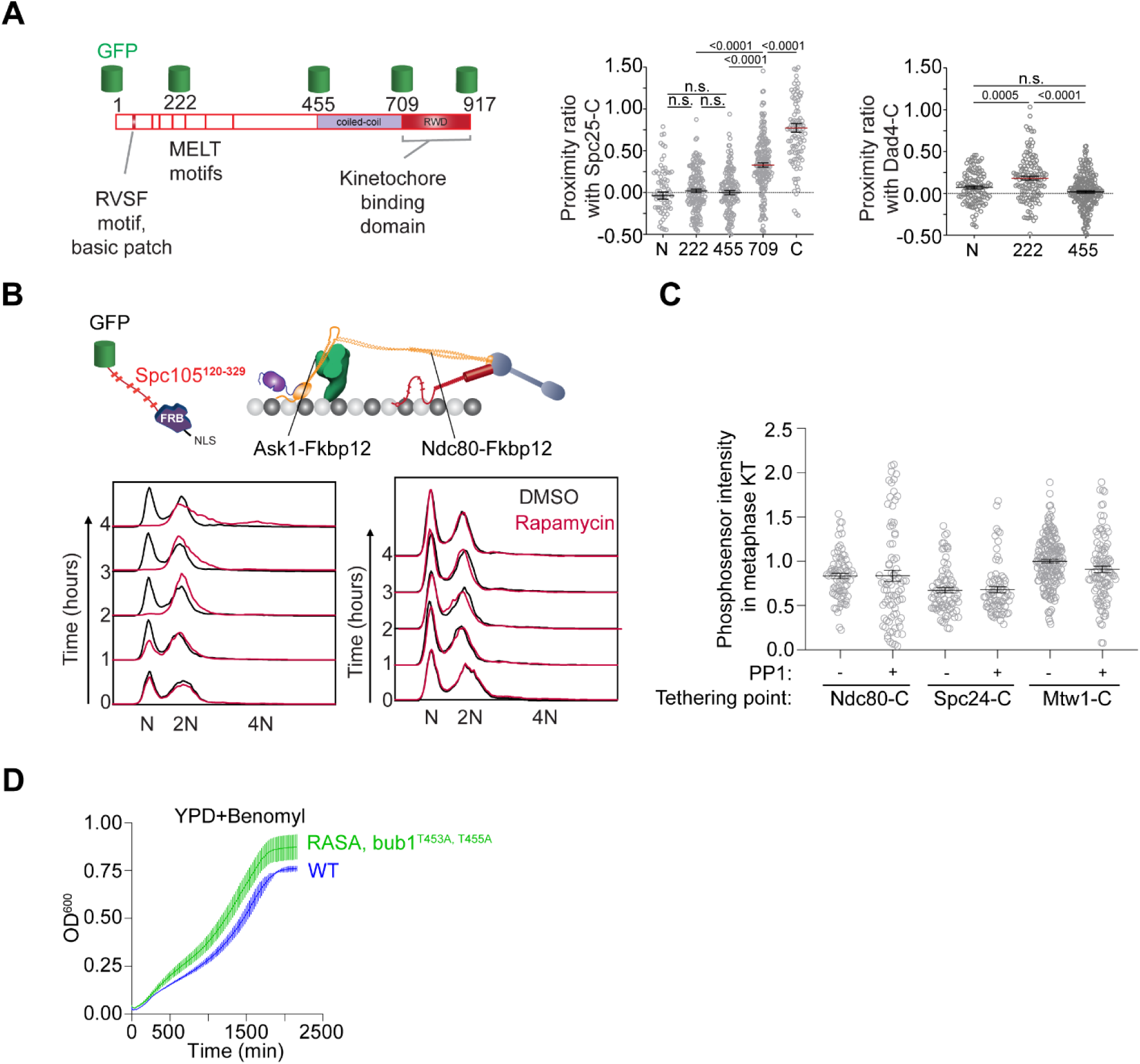
The distribution of the Spc105 phosphodomain in bioriented kinetochores and the position-dependent effects of tethering an additional phosphodomain within the kinetochore. **A** The assessment of the nanoscale distribution of the unstructured phosphodomain of Spc105 using FRET. Top: Schematic displays the GFP-insertion positions within Spc105. Bottom: Scatter plots show the proximity ratio (which is proportional to FRET) of donor GFP fused at the indicated residue in Spc105 with Spc25 C-terminus (left) and Dad4 C-terminus (right). (mean+s.e.m. n= 74, 165, 142, 177 and 103 respectively for FRET involving Spc25-mCherry, accumulated from at least three repeats. n= 114, 134 and 224 for N, 222 and 455 respectively from three repeats for FRET involving Dad4-mCherry, accumulated from at least two repeats). n.s. Not significant. **B** Flow cytometry-based quantification of the DNA content in cells treated with rapamycin (prior to sample collection at 0 minutes) to tether the phosphosensor at the indicated position. Representative histogram from two trials is presented here. **C** Quantification of amount of Spc105 phosphodomain tethered to bioriented kinetochore clusters (mean+s.e.m. n=74 and 80 for Ndc80-C, 78 and 76 for Spc24-C, 102 and 173 for Mtw1-C, pooled from at least two experimental replicates). **D** Quantification of population growth for the indicated strains in presence of benomyl (30µg/ml) (mean± s.e.m., n= at least 3).

**Figure S5 (related to Figure 5 and 6).**
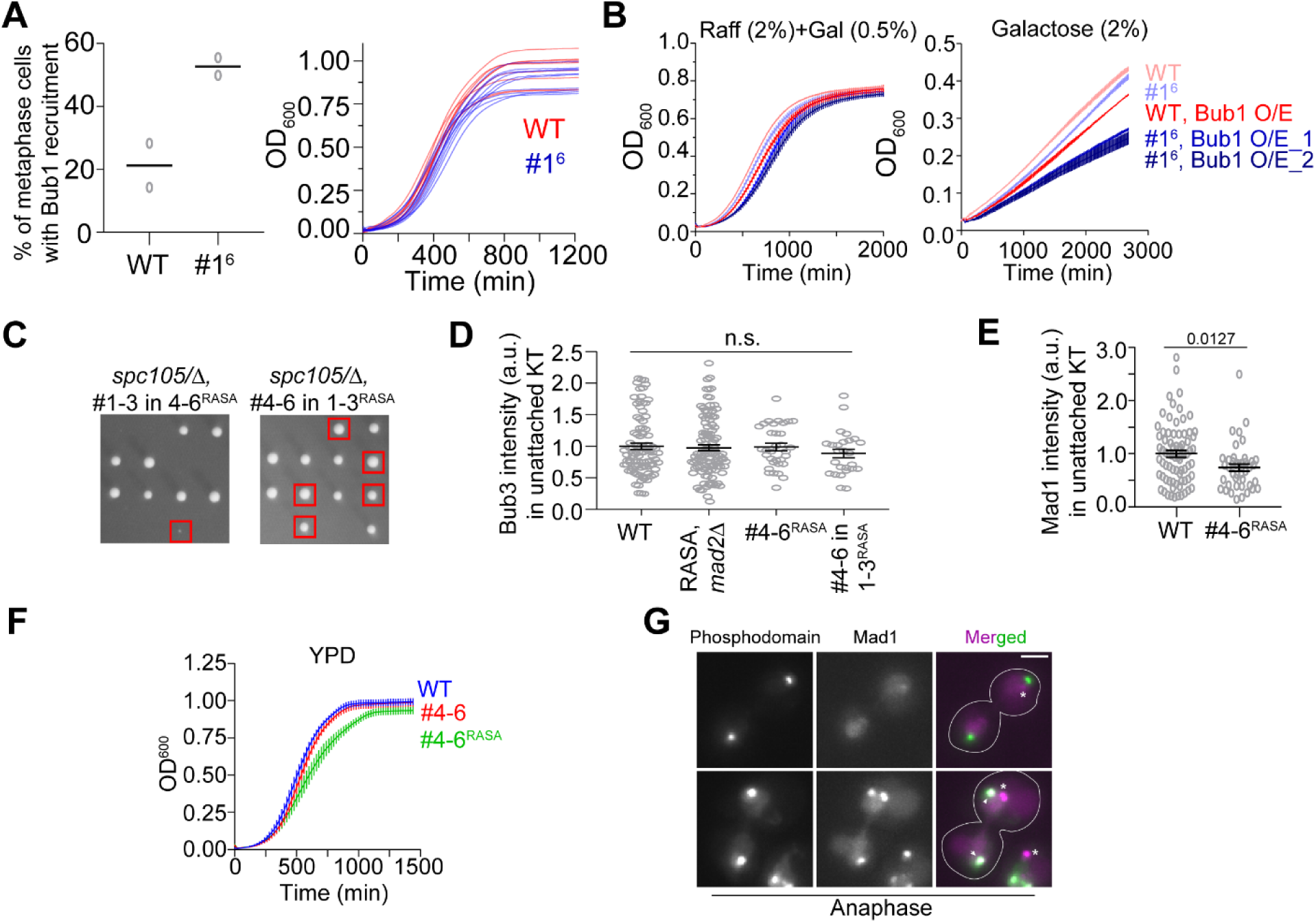
The influence of the affinity of MELT motifs to Bub3-Bub1 expression on the responsiveness of the SAC to silencing mechanisms. **A** Scatter plot showing percentage of cells with bioriented kinetochores with Bub1 localization in the indicated strains. (n=211 and 74 for WT and MELT1^6^ respectively, horizontal line indicates mean value). **B** Growth curves of the indicated strains in presence of raffinose+galactose (left) and in presence of galactose (right) as measured by absorbance at 600nm (mean±s.e.m. shown for each time point from 3 replicates). **C** Tetrad dissection analysis of the indicated strains. Red squares mark the tetrads of the desirable genotype. The dissections were repeated with two biological replicates. **D** Quantification of Bub3-mCherry intensity values of the indicated strains. (mean+s.e.m. n=92, 107, 33 and 26 for WT, *spc105^RASA, #4-6^*, *mad2*Δ, *spc^105RASA, #4-6^* and *spc105^RASA, #4-6in1-3^* respectively, pooled from two repeats of the experiment). n.s. Not significant. **E** Quantitative comparison of the amount of Mad1-mCherry recruited by unattached kinetochores in *spc105^RASA, #4-6^* cells treated with nocodazole. n=74 and 42 for WT and *spc105^RASA, #4-6^* respectively, pooled from two experimental repeats. **F** Left: Growth curve shows quantification of the evolution of cell density of the indicated strains in YPD. **G** Representative micrographs display Mad1-mCherry localization relative to kinetochores of anaphase cells in the indicated strains after treatment with rapamycin. The Mad1-mCherry puncta marked with an asterisk result from the deletion of the nuclear pore protein Nup60. These puncta are not associated with kinetochores. The ones co-localized with kinetochores are marked with arrowheads.

## References

Aravamudhan, P., Chen, R., Roy, B., Sim, J., and Joglekar, A.P. (2016). Dual mechanisms regulate the recruitment of spindle assembly checkpoint proteins to the budding yeast kinetochore. Mol Biol Cell 27, 3405–3417.

Aravamudhan, P., Felzer-Kim, I., Gurunathan, K., and Joglekar, A.P. (2014). Assembling the protein architecture of the budding yeast kinetochore-microtubule attachment using FRET. Curr Biol 24, 1437–1446.

Aravamudhan, P., Goldfarb, A.A., and Joglekar, A.P. (2015). The kinetochore encodes a mechanical switch to disrupt spindle assembly checkpoint signalling. Nat Cell Biol 17, 868–879.

Chen, C., Whitney, I.P., Banerjee, A., Sacristan, C., Sekhri, P., Kern, D.M., Fontan, A., Kops, G., Tyson, J.J., Cheeseman, I.M., et al. (2019). Ectopic Activation of the Spindle Assembly Checkpoint Signaling Cascade Reveals Its Biochemical Design. Curr Biol 29, 104–119 e110.

Duffy, S., and Hieter, P. (2018). The Chromosome Transmission Fidelity Assay for Measuring Chromosome Loss in Yeast. Methods in molecular biology 1672, 11–19.

Faesen, A.C., Thanasoula, M., Maffini, S., Breit, C., Muller, F., van Gerwen, S., Bange, T., and Musacchio, A. (2017). Basis of catalytic assembly of the mitotic checkpoint complex. Nature.

Gillett, E.S., Espelin, C.W., and Sorger, P.K. (2004). Spindle checkpoint proteins and chromosome– microtubule attachment in budding yeast. The Journal of Cell Biology 164, 535–546.

Gupta, A., Evans, R.K., Koch, L.B., Littleton, A.J., and Biggins, S. (2018). Purification of kinetochores from the budding yeast Saccharomyces cerevisiae. Methods in cell biology 144, 349–370.

Hieter, P., Mann, C., Snyder, M., and Davis, R.W. (1985). Mitotic stability of yeast chromosomes: a colony color assay that measures nondisjunction and chromosome loss. Cell 40, 381–392.

Hiruma, Y., Sacristan, C., Pachis, S.T., Adamopoulos, A., Kuijt, T., Ubbink, M., von Castelmur, E., Perrakis, A., and Kops, G.J. (2015). CELL DIVISION CYCLE. Competition between MPS1 and microtubules at kinetochores regulates spindle checkpoint signaling. Science 348, 1264–1267.

Hung, C.W., Martinez-Marquez, J.Y., Javed, F.T., and Duncan, M.C. (2018). A simple and inexpensive quantitative technique for determining chemical sensitivity in Saccharomyces cerevisiae. Sci Rep 8, 11919.

Janke, C., Magiera, M.M., Rathfelder, N., Taxis, C., Reber, S., Maekawa, H., Moreno-Borchart, A., Doenges, G., Schwob, E., Schiebel, E., et al. (2004). A versatile toolbox for PCR-based tagging of yeast genes: new fluorescent proteins, more markers and promoter substitution cassettes. Yeast 21, 947–962.

Ji, Z., Gao, H., Jia, L., Li, B., and Yu, H. (2017). A sequential multi-target Mps1 phosphorylation cascade promotes spindle checkpoint signaling. Elife 6.

Ji, Z., Gao, H., and Yu, H. (2015). CELL DIVISION CYCLE. Kinetochore attachment sensed by competitive Mps1 and microtubule binding to Ndc80C. Science 348, 1260–1264.

Jin, F., Liu, H., Li, P., Yu, H.G., and Wang, Y. (2012). Loss of function of the Cik1/Kar3 motor complex results in chromosomes with syntelic attachment that are sensed by the tension checkpoint. PLoS Genet 8, e1002492.

Joglekar, A., Chen, R., and Lawrimore, J. (2013). A Sensitized Emission Based Calibration of FRET Efficiency for Probing the Architecture of Macromolecular Machines. Cell Mol Bioeng 6, 369–382.

Joglekar, A.P., Bloom, K., and Salmon, E.D. (2009). In vivo protein architecture of the eukaryotic kinetochore with nanometer scale accuracy. Curr Biol 19, 694–699.

Joglekar, A.P., Bouck, D.C., Molk, J.N., Bloom, K.S., and Salmon, E.D. (2006). Molecular architecture of a kinetochore–microtubule attachment site. Nature Cell Biology 8, 581.

Jürg Bähler, Jian-Qiu Wu, Mark S. Longtine, Nirav G. Shah, Amos Mckenzie III, Alexander B. Steever, Achim Wach, Peter Philippsen, and Pringle, J.R. (1998). Heterologous modules for efficient and versatile PCR-based gene targeting in Schizosaccharomyces pombe. YEAST 14, 943–951.

Kawashima, S.A., Yamagishi, Y., Honda, T., Ishiguro, K.-i., and Watanabe, Y. (2010a). Phosphorylation of H2A by Bub1 Prevents Chromosomal Instability Through Localizing Shugoshin. Science 327, 172–177.

Kawashima, S.A., Yamagishi, Y., Honda, T., Ishiguro, K., and Watanabe, Y. (2010b). Phosphorylation of H2A by Bub1 prevents chromosomal instability through localizing shugoshin. Science 327, 172–177.

Kemmler, S., Stach, M., Knapp, M., Ortiz, J., Pfannstiel, J., Ruppert, T., and Lechner, J. (2009). Mimicking Ndc80 phosphorylation triggers spindle assembly checkpoint signalling. EMBO J 28, 1099–1110.

Koch, L.B., Opoku, K.N., Deng, Y., Barber, A., Littleton, A.J., London, N., Biggins, S., and Asbury, C.L. (2019). Autophosphorylation is sufficient to release Mps1 kinase from native kinetochores. Proc Natl Acad Sci USA 116, 17355–17360.

Krenn, V., Overlack, K., Primorac, I., van Gerwen, S., and Musacchio, A. (2014). KI motifs of human Knl1 enhance assembly of comprehensive spindle checkpoint complexes around MELT repeats. Curr Biol 24, 29–39.

Liu, D., Vleugel, M., Backer, C.B., Hori, T., Fukagawa, T., Cheeseman, I.M., and Lampson, M.A. (2010). Regulated targeting of protein phosphatase 1 to the outer kinetochore by KNL1 opposes Aurora B kinase. J Cell Biol 188, 809–820.

London, N., and Biggins, S. (2014). Mad1 kinetochore recruitment by Mps1-mediated phosphorylation of Bub1 signals the spindle checkpoint. Genes Dev 28, 140–152.

London, N., Ceto, S., Ranish, J.A., and Biggins, S. (2012). Phosphoregulation of Spc105 by Mps1 and PP1 regulates Bub1 localization to kinetochores. Curr Biol 22, 900–906.

Meadows, J.C., Shepperd, L.A., Vanoosthuyse, V., Lancaster, T.C., Sochaj, A.M., Buttrick, G.J., Hardwick, K.G., and Millar, J.B. (2011a). Spindle checkpoint silencing requires association of PP1 to both Spc7 and kinesin-8 motors. Dev Cell 20, 739–750.

Meadows, John C., Shepperd, Lindsey A., Vanoosthuyse, V., Lancaster, Theresa C., Sochaj, Alicja M., Buttrick, Graham J., Hardwick, Kevin G., and Millar, Jonathan B.A. (2011b). Spindle Checkpoint Silencing Requires Association of PP1 to Both Spc7 and Kinesin-8 Motors. Developmental Cell 20, 739–750.

Musacchio, A. (2015). The Molecular Biology of Spindle Assembly Checkpoint Signaling Dynamics. Curr Biol 25, R1002–1018.

Overlack, K., Primorac, I., Vleugel, M., Krenn, V., Maffini, S., Hoffmann, I., Kops, G.J., and Musacchio, A. (2015). A molecular basis for the differential roles of Bub1 and BubR1 in the spindle assembly checkpoint. Elife 4, e05269.

Peplowska, K., Wallek, A.U., and Storchova, Z. (2014). Sgo1 regulates both condensin and Ipl1/Aurora B to promote chromosome biorientation. PLoS Genet 10, e1004411.

Primorac, I., Weir, J.R., Chiroli, E., Gross, F., Hoffmann, I., van Gerwen, S., Ciliberto, A., and Musacchio, A. (2013). Bub3 reads phosphorylated MELT repeats to promote spindle assembly checkpoint signaling. Elife 2, e01030.

Qian, J., Garcia-Gimeno, M.A., Beullens, M., Manzione, M.G., Van der Hoeven, G., Igual, J.C., Heredia, M., Sanz, P., Gelens, L., and Bollen, M. (2017). An Attachment-Independent Biochemical Timer of the Spindle Assembly Checkpoint. Mol Cell 68, 715–730 e715.

Rosenberg, J.S., Cross, F.R., and Funabiki, H. (2011). KNL1/Spc105 recruits PP1 to silence the spindle assembly checkpoint. Curr Biol 21, 942–947.

Roy, B., Verma, V., Sim, J., Fontan, A., and Joglekar, A.P. (2019). Delineating the contribution of Spc105-bound PP1 to spindle checkpoint silencing and kinetochore microtubule attachment regulation. J Cell Biol.

Scott, R.J., Lusk, C.P., Dilworth, D.J., Aitchison, J.D., and Wozniak, R.W. (2005). Interactions between Mad1p and the nuclear transport machinery in the yeast Saccharomyces cerevisiae. Mol Biol Cell 16, 4362–4374.

Tanaka, K., Mukae, N., Dewar, H., van Breugel, M., James, E.K., Prescott, A.R., Antony, C., and Tanaka, T.U. (2005). Molecular mechanisms of kinetochore capture by spindle microtubules. Nature 434, 987–994.

Tipton, A.R., Ji, W., Sturt-Gillespie, B., Bekier, M.E., 2nd, Wang, K., Taylor, W.R., and Liu, S.T. (2013). Monopolar spindle 1 (MPS1) kinase promotes production of closed MAD2 (C-MAD2) conformer and assembly of the mitotic checkpoint complex. J Biol Chem 288, 35149–35158.

Tromer, E., Snel, B., and Kops, G.J. (2015). Widespread Recurrent Patterns of Rapid Repeat Evolution in the Kinetochore Scaffold KNL1. Genome Biol Evol 7, 2383–2393.

Vleugel, M., Omerzu, M., Groenewold, V., Hadders, M.A., Lens, S.M., and Kops, G.J. (2015). Sequential multisite phospho-regulation of KNL1-BUB3 interfaces at mitotic kinetochores. Mol Cell 57, 824–835.

Vleugel, M., Tromer, E., Omerzu, M., Groenewold, V., Nijenhuis, W., Snel, B., and Kops, G.J. (2013). Arrayed BUB recruitment modules in the kinetochore scaffold KNL1 promote accurate chromosome segregation. J Cell Biol 203, 943–955.

Warren, C.D., Brady, D.M., Johnston, R.C., Hanna, J.S., Hardwick, K.G., and Spencer, F.A. (2002). Distinct chromosome segregation roles for spindle checkpoint proteins. Mol Biol Cell 13, 3029–3041.

Zhang, G., Lischetti, T., and Nilsson, J. (2014). A minimal number of MELT repeats supports all the functions of KNL1 in chromosome segregation. J Cell Sci 127, 871–884.

